# GIRAF 1.0: A unified global framework to anticipate plant pest invasions

**DOI:** 10.64898/2026.04.23.720440

**Authors:** Aaron I. Plex Sulá, Ozgur Batuman, Gilles Cellier, Nicholas S. Dufault, Berea A. Etherton, Amanda Hodges, Tiffany M. Lowe-Power, John D. McVay, Cory Penca, Kyle Schroeder, Eleni Stilian, Piotr Suder, Yu Takeuchi, Henri E. Z. Tonnang, Ying Wang, Karen A. Garrett

**Affiliations:** Plant Pathology Department, University of Florida, Gainesville, FL, USA 32611; Emerging Pathogens Institute, University of Florida, Gainesville, FL, USA 32611; Global Food Systems Institute, University of Florida, Gainesville, FL, USA 32611; Southwest Florida Research and Education Center (SWFREC), University of Florida, Immokalee, FL, USA 34142; ANSES, Plant Health Laboratory, Saint-Pierre, Reunion Island, France 97410; Department of Entomology and Nematology, University of Florida, Gainesville, FL, USA 32611; Department of Plant Pathology, University of California Davis, Davis, CA, USA 95616; Florida Department of Agriculture and Consumer Services (FDACS) Division of Plant Industry, Gainesville, FL, USA 32608; USDA APHIS PPQ S&T 13601 Old Cutler Road, Miami, Florida, USA 33158; Department of Statistical Science, Duke University, Durham, NC, USA 27705; Center for Integrated Pest Management, North Carolina State University, Raleigh, NC, USA 27606; International Institute of Tropical Agriculture (IITA), Ibadan, Nigeria 200001; School of Agricultural, Earth, and Environmental Sciences, University of KwaZulu-Natal, Pietermaritzburg, South Africa 3209

## Abstract

Plant pests threaten 10-40% of global food production, resulting in $55-220 billion in annual economic losses. Despite these escalating risks, biosecurity remains largely reactive, lacking anticipatory frameworks that integrate pest-specific drivers governing transboundary spread. We present GIRAF 1.0 (Global Invasion Risk Assessment Framework), the first quantitative, data-driven system that unifies pest-specific multi-host landscapes, abiotic suitability, and global trade networks with international phytosanitary policies. We applied GIRAF to four globally devastating pests – ranging from viral to insect taxa – to reconstruct a century of transcontinental spread and generate the first multiscale atlases of future invasion potential. GIRAF reveals that 22-37% of Earth’s land surface can contain host communities that largely overlap with environmentally suitable hotspots. Over 115 countries are highly vulnerable to trade-mediated pest introductions despite adopted phytosanitary policies. GIRAF provides a foundation for proactive surveillance and pandemic preparedness, offering a scalable path for transnational biosecurity agencies and global food industries.

## Introduction

Plant ecosystems are currently undermined by a 10-40% loss in crop yield caused by new and re-emerging diseases and pests globally^1^. Unchecked outbreaks of plant diseases and transboundary pests decimate global food baskets, disrupt international markets, and threaten natural ecosystem functions^2^. This global crisis in pest invasions has accelerated due to the combined impacts of intensified commodity trade, human mobility, cropland expansion, and climate change^3-6^.

National plant protection strategies are vital to achieving the UN Sustainable Development Goals (SDGs)^7^, but remain largely reactive to transcontinental pest outbreaks and pandemics in the 21^st^ century^6,8^. Global proactive biosecurity – our collective capacity to prevent destructive pests from expanding and persisting in new biogeographical regions – requires integrated, spatially explicit, species-specific risk assessments^3,7,9^. This anticipatory approach is increasingly needed to identify the most likely locations for introduction, establishment, and spread across pest species^9-12^, informing a One Biosecurity policy framework^7^.

Nevertheless, anticipating the actual global spread of invasive pests involves a high degree of uncertainty, including unpredictable stochastic processes. Currently, geographic risk analysis of pest invasions often relies on qualitative expert opinion, which may be the only option for some species, or climate-based risk projection^13^. Hundreds of quick risk assessments are available for specific nations^9^, but they represent a small fraction of a growing number of pests affecting plants^8^.

Understanding of individual drivers of pest invasions has substantially increased in the last three decades^13-15^. However, we lack a unified, data-driven framework that integrates multiple species-specific drivers across global and local scales. This challenge is exacerbated by persisting data gaps in high-resolution host distribution, trade flows of high-risk commodities, and species-specific environmental requirements^16,17^. To bridge this gap, invasion science must build on limited geospatial data with targeted yet generalizable, data-driven principles to explicitly quantify invasion risks^18^.

Here, we present GIRAF 1.0 (Global Invasion Risk Assessment Framework): a quantitative, use-inspired system for evaluating potential scenarios of the geographic spread of invasive pests (Fig. 1). GIRAF offers a data-driven foundation that complements expert-driven assessments. GIRAF integrates for the first time four fundamental drivers of pest invasions and pandemic potential in global plant ecosystems: (i) international trade of high-risk commodities^19,20^, (ii) cropland accessibility to ports and urban entry points, (iii) host landscape connectivity determining local spread potential^21^, and (iv) environmental suitability based on climate and edaphic constraints^9^ (Fig. 1). By identifying the most likely paths of spread, GIRAF maps candidate priority locations for global surveillance planning^5,22^.

**Fig. 1.**
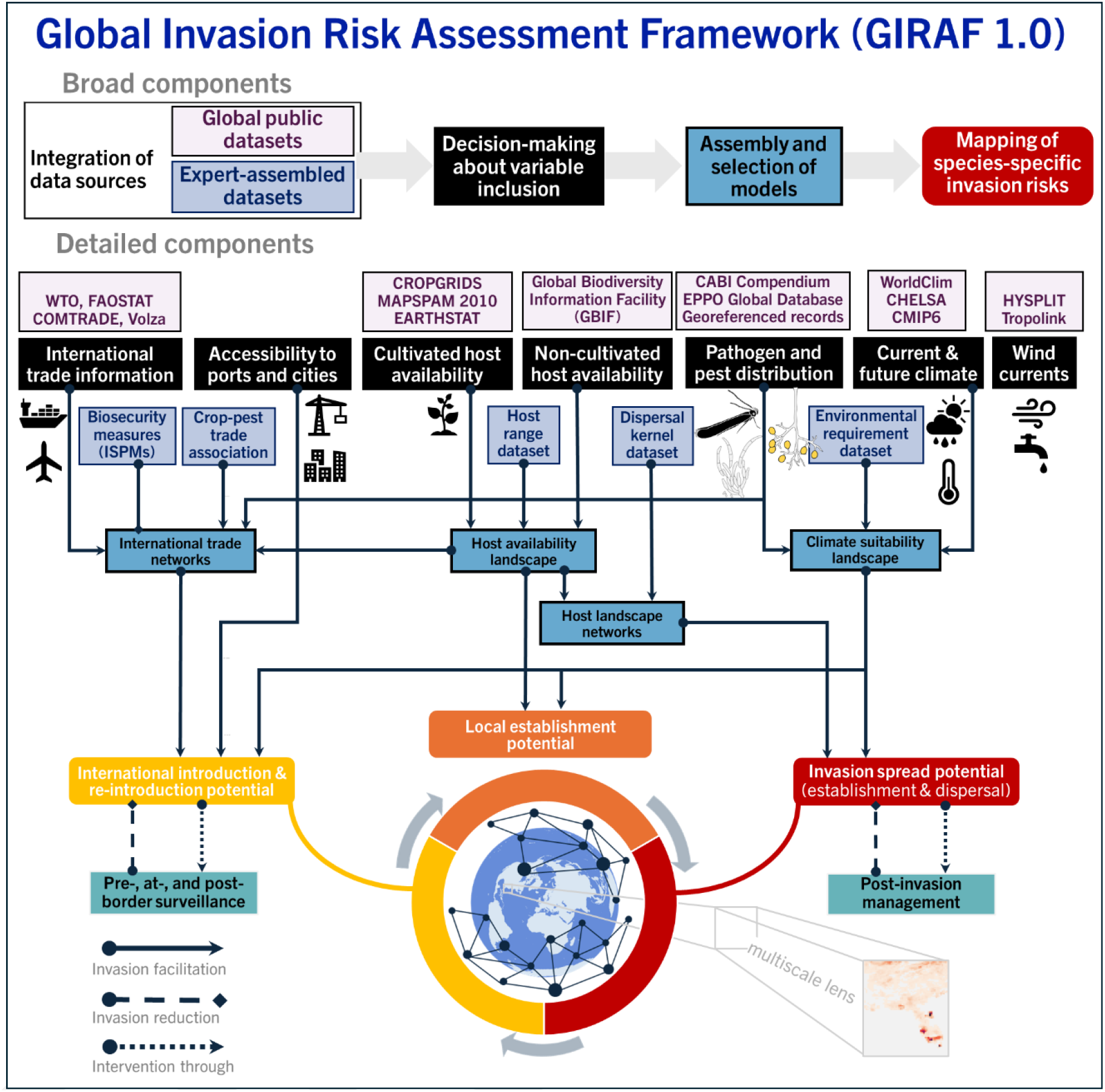
GIRAF 1.0, a new global framework for assessing invasion potential. GIRAF 1.0 integrates four fundamental drivers of invasive pest spread: environmental suitability, host connectivity, international trade (e.g., seed exchange), and local transportation (e.g., access to cities). GIRAF comprises four general modeling processes: integration of data sources (public datasets and expert-assembled datasets), decision-making about variable inclusion, model assembly and selection, and geographic predictions of invasion risk. In GIRAF, expert evaluation is needed throughout the modeling process, from data input to model selection, parameter choices, and the importance of each geographic driver. GIRAF’s primary goal is to produce species-specific invasion risk maps for (pro)active surveillance and risk mitigation over a contemporary time horizon. GIRAF is applied across geographic scales, from global to local.

To demonstrate GIRAF’s taxonomic and geographic generalizability, we assess the global risks posed by economically and globally devastating pests representing insect, bacterial, viral, and viroid taxa: the South American tomato leafminer (*Phthorimaea absoluta*), *Ralstonia solanacearum* phylotype IIB sequevar 1 (RSIIB-1; formerly designated as race 3 biovar 2), tomato brown rugose fruit virus (*Tobamovirus fructirugosum*; ToBRFV), and potato spindle tuber viroid (*Pospiviroid fusituberis*; PSTVd). These transboundary pests threaten the US$334 billion solanaceous crop industry (peppers, potatoes, and tomatoes)^23^, ornamental industries, and natural plant ecosystems, cornerstones of global food security and livelihoods.

By reconstructing a century of transcontinental range expansions of these pests, we identify high-probability invasion scenarios in global plant ecosystems (Fig. S1-2). A variety of generic models for invasion dynamics exists^3,20^. GIRAF augments these generic models by accounting for the specific ecological niches of each pest, defined by host distribution, dispersal pathways, and environmental drivers (Note S1-2). Lastly, we show GIRAF’s multi-scale lens by downscaling our global analysis to quantify local invasion potential in the Southeastern United States, providing a foundational tool for proactive transnational biosecurity.

## Results

### Introduction vulnerabilities in global trade networks

While local sentinel surveillance and ongoing management need to continue actively in regions where pests are already present (Fig. S1-2), proactive surveillance can target potential accidental pest movement among countries through international trade (Fig. 2). While previous risk assessments often rely on climate suitability alone – such as climate-based assessment for *P. absoluta*^*24*^ – GIRAF 1.0 integrates historical trade flows of pest-associated commodities to reveal previously unquantified introduction hotspots.

**Fig. 2.**
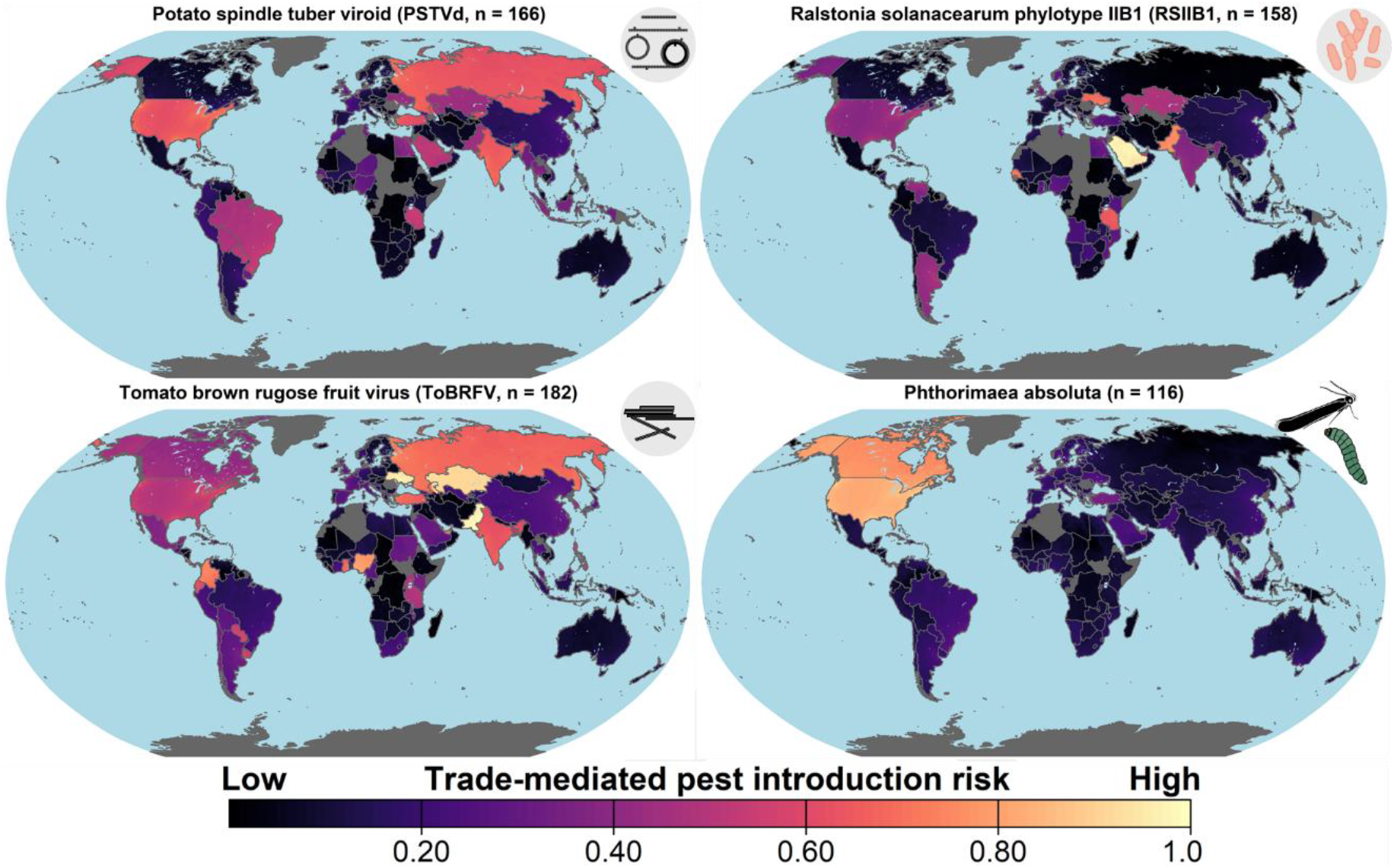
Cumulative vulnerability to (re)introduction of four invasive plant pests. This vulnerability analysis considers international pest-specific commodity pathways, pest distribution at the country level, major crop host availability at the country level, and accessibility to ports. n is the number of importing countries with a possible introduction vulnerability through international trade of agricultural commodities. Countries in grey have no reports of international trade of these commodities.

We found that over 115 countries are potential destinations for at least one of the pests. Our analysis reveals that major producers of solanaceous crops serve as critical nodes in global trade networks and are highly vulnerable to the potential introduction of all four target pests. Highly vulnerable countries include the USA, Canada, Bahrain, Kuwait, and UAE for *P. absoluta*; the USA, Niger, Saudi Arabia, and Egypt for PSTVd; Pakistan, Tanzania, Saudi Arabia, and Senegal for RSIIB-1; and Ukraine, the USA, Canada, and Kazakhstan for ToBRFV (Fig. 2).

Phytosanitary measures specific to each pest are implemented by a few countries (Data S2-7). Each pest has a global yet scattered distribution (Fig. S1-2). Many countries exporting agricultural commodities may serve as possible pest source pools (Fig. 3). If these historical patterns in international trade of high-risk commodities persist, most solanaceous crop-growing countries will remain highly vulnerable to potential trade-mediated introductions. International trade in seed, planting material, and food products has reportedly played a key role in the rapid global spread of these pests (Data S10). These global patterns render long-term prevention of pest introductions through safe international trade increasingly challenging.

**Fig. 3.**
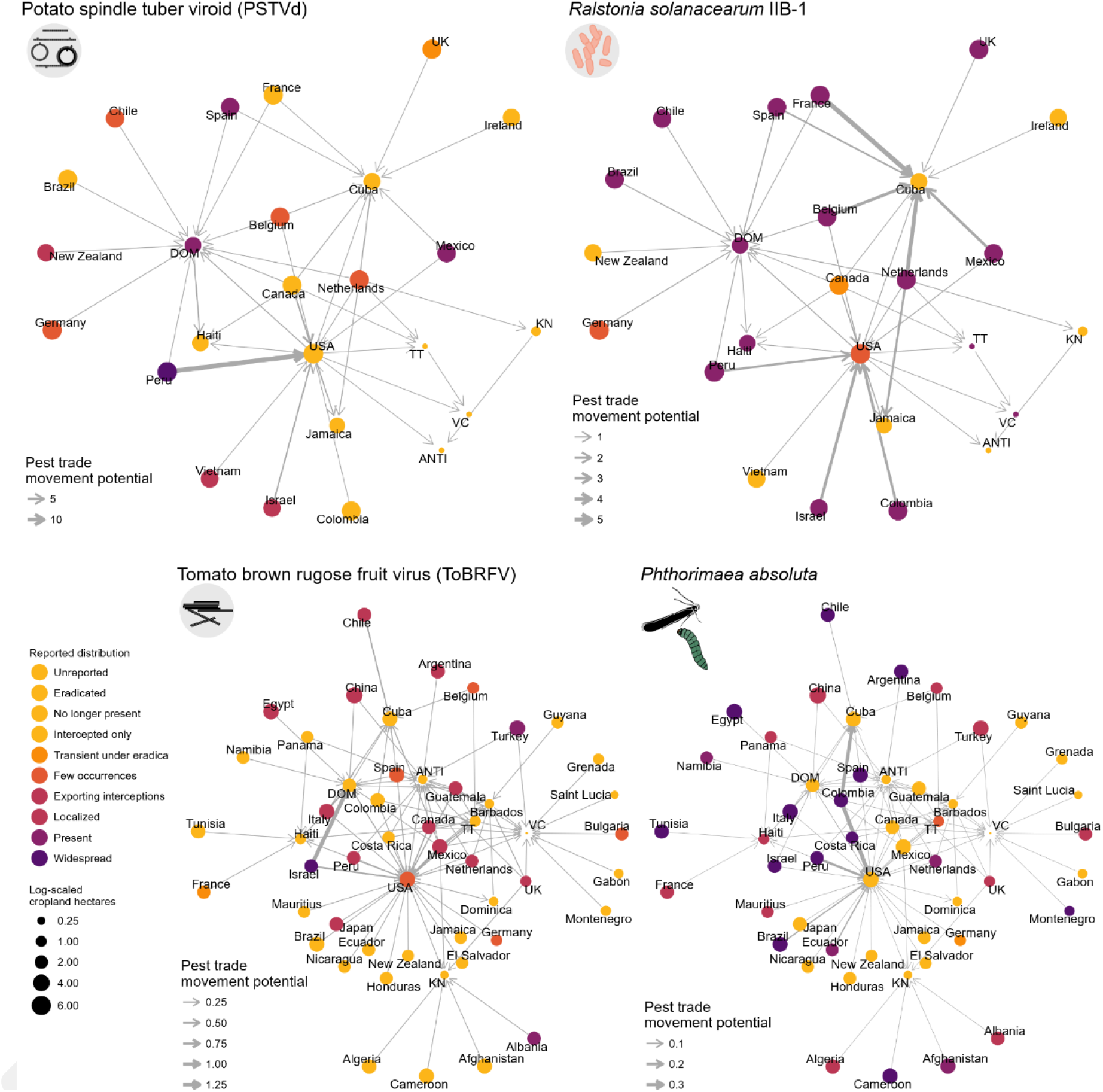
Potential geographic spread routes of pathogens and pests through international trade networks linked to the Caribbean region and United States. Countries closer to the center of the trade networks have higher vulnerability to pathogen or pest introduction via trade of high-risk crop commodities. Node size indicates the availability of crop species that are hosts of the pest. Pathogen or pest invasion potential and trade movement potential are relative rankings to compare invasion potential in a species-specific trade network (not for comparison between species networks). Abbreviations: ANTI – Antigua and Barbuda, DOM – Dominican Republic, KN – Saint Kitts and Nevis, TT – Trinidad and Tobago, VC – Saint Vincent and the Grenadines.

GIRAF can also support assessment of the potential for pest introduction in specific geographic regions. For example, for proactive surveillance prioritization in the USA and the Caribbean region, GIRAF identified key regional hubs in trade networks that are more vulnerable to the (re)introduction of each pest (Fig. 3). These regional hubs consist of host-growing countries that import pest-associated crop commodities from many regions where the target pest is present. The USA, the Dominican Republic, and Cuba serve as regional trade hubs, experiencing high introduction vulnerability to each pest. Likewise, Saint Vincent, the Grenadines, Antigua, and Barbuda may act as possible trade hubs vulnerable to *P. absoluta* or ToBRFV (re)introduction.

GIRAF also distinguished spatially explicit, potential movement pathways for each target pest (Fig. 3). For example, while Cuba’s potato seed imports from nine countries represent a high vulnerability to RSIIB-1 introduction, the same trade network poses a substantially lower vulnerability to PSTVd (Fig. 3). Similarly, the USA imported tomato fruit and seed from 27 countries: these trade connections provide a possible movement pathway for *P. absoluta* from South America, but low movement potential for ToBRFV. This capacity to distinguish risks across taxonomically diverse pests within the same trade framework allows National Plant Protection Organizations (NPPOs) to target mobilization of limited resources.

### Invasion vulnerabilities based on environmental suitability

Once a pest bypasses trade-based barriers, national early warning systems need to determine the degree and extent to which the local environment is suitable for establishment. GIRAF identifies suitable environments for pest invasions based on climatic, edaphic, and hydrological conditions. For each pest, GIRAF inferred environmental suitability from a machine-learning ensemble trained on the most comprehensive compilation of geospatially distinct outbreak observations over the last century.

This environment-based modeling approach predicted the currently reported georeferenced range of each target pest (Fig. 4), where the average accuracy ranged from 61% for PSTVd based on logistic regression to 96% for ToBRFV based on Maxent (Fig. S3). GIRAF provides spatial predictions beyond reported geographical ranges of each target pest, indicating which locations might be environmentally suitable for pest development under contemporary conditions.

**Fig. 4.**
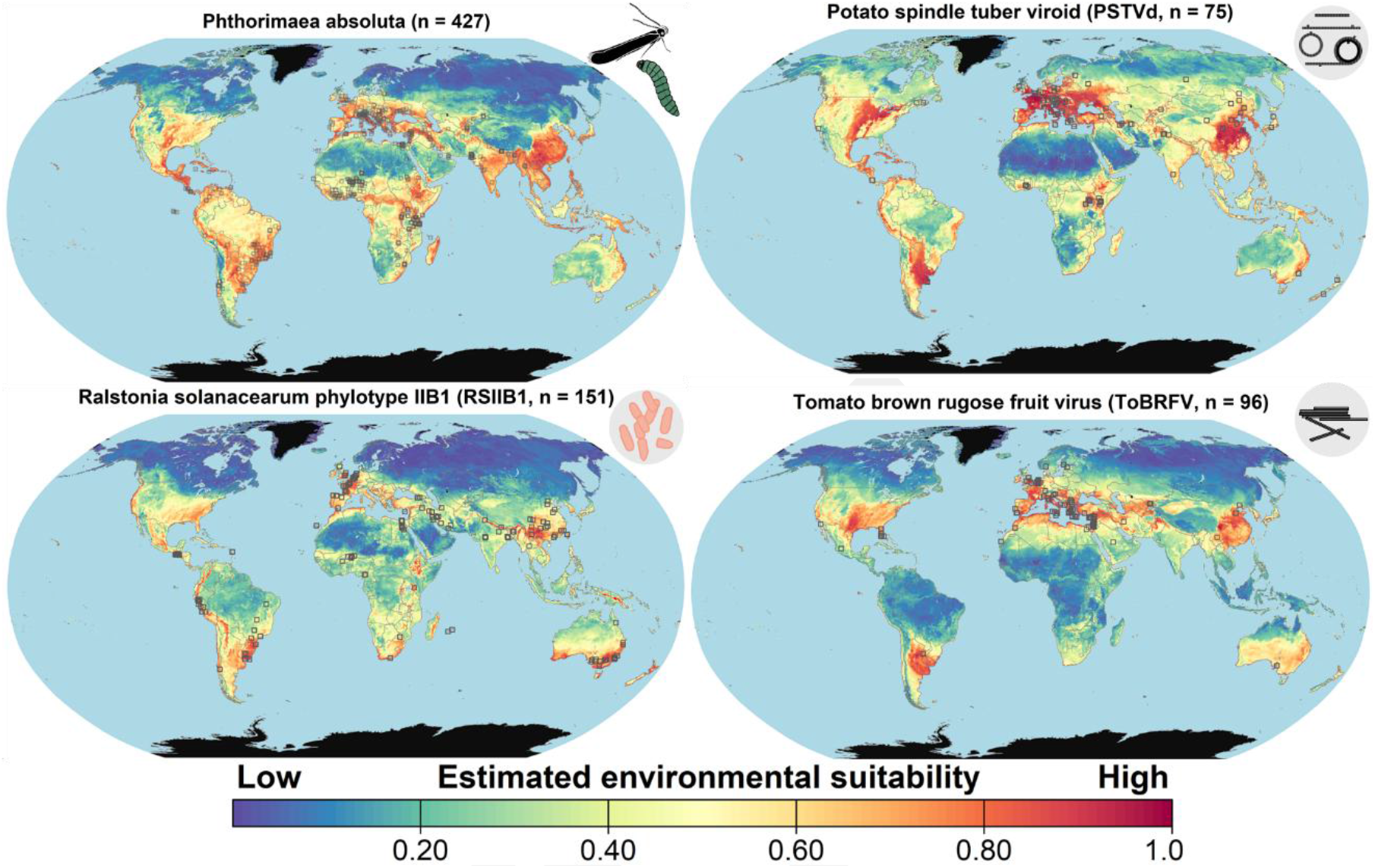
Global vulnerability hotspots for four invasive pests based on an ensemble of environment-based machine-learning models. Sample size (n) is the number of historical georeferenced pest outbreak observations (black square outlines) used to train individual species distribution models. Black pixels are locations the ensemble model predicted to be environmentally unsuitable.

Environmentally suitable hotspots – regions with >50% relative likelihood of invasion for each of the four pests – include a large area in China, and Eastern Australia (Fig. 4), where each pest has been reported (Fig. S1). Likewise, highly suitable environments are available throughout regions in Europe, where each pest is reportedly present in different countries. The Southeastern USA and the US Pacific Coast are probably environmentally suitable areas for each pest, even though these pests are currently not reported in these regions.

Other potentially suitable hotspots are specific to each target pest (Fig. 4). For example, Northern and central Argentina, the Chaco in Paraguay, the tropical Andes in Bolivia, Peru, Ecuador, Colombia, and Venezuela, most of Central America, and the Atlantic coastal regions in Mexico are likely suitable for PSTVd. Despite the lack of georeferenced occurrences in these regions, our analysis reveals within-country patterns of environmental suitability for the first time. These findings are, though, validated by pest occurrence data collected nationally (Fig. S1). Reports of RSIIB-1 are scattered in the Mediterranean basin, but our model indicates high environmental suitability in these countries. GIRAF identifies these environmentally suitable hotspots where proactive surveillance programs could be targeted to mitigate the potential risks from range expansion.

### Invasion vulnerabilities based on multi-host landscape connectivity

A critical limitation in current biosecurity is the reliance on maps of single host species (usually a crop), which ignores the likely role of non-cultivated hosts in pest spread. GIRAF 1.0 addresses this gap by mapping the global distribution of multiple hosts of a target pest using publicly accessible databases. Furthermore, GIRAF provides connectivity assessments for crops and non-cultivated hosts of a pest species. Here, host connectivity refers to the relative likelihood of pest spread between host locations if the pest reaches a target location within the host landscape, all else being equal. GIRAF quantifies potential host connectivity based on host availability (structural connectivity) and a highly likely range of pest dispersal parameters. Mapping multi-species connectivity is a first approximation for the potential spread of each target pest in realistic, heterogeneous host landscapes. This approach identified not only potential structural geographic barriers where a host is unreported but also spatially contiguous host areas and fragmented host habitats.

Host availability strongly correlates with mean host connectivity (Spearsman’s ***r***_***s***_ = 0.62 for PSTVd, 0.81 for RSIIB-1, 0.65 for ToBRFV, and 0.80 for *P. absoluta*, ***p*** < 2.2e-16), so high-density host communities commonly have high functional connectivity (Fig. S4-9). This pattern is supported by small differences in ranks for locations with high host connectivity and high host availability (Fig. S5). For example, Europe, Southern Asia, and China have a homogeneously dense host landscape that is likely to facilitate the local spread of each pest (Fig. 5). A homogeneously dense host landscape for *P. absoluta* is also available throughout Central and North America (Fig. 5).

**Fig. 5.**
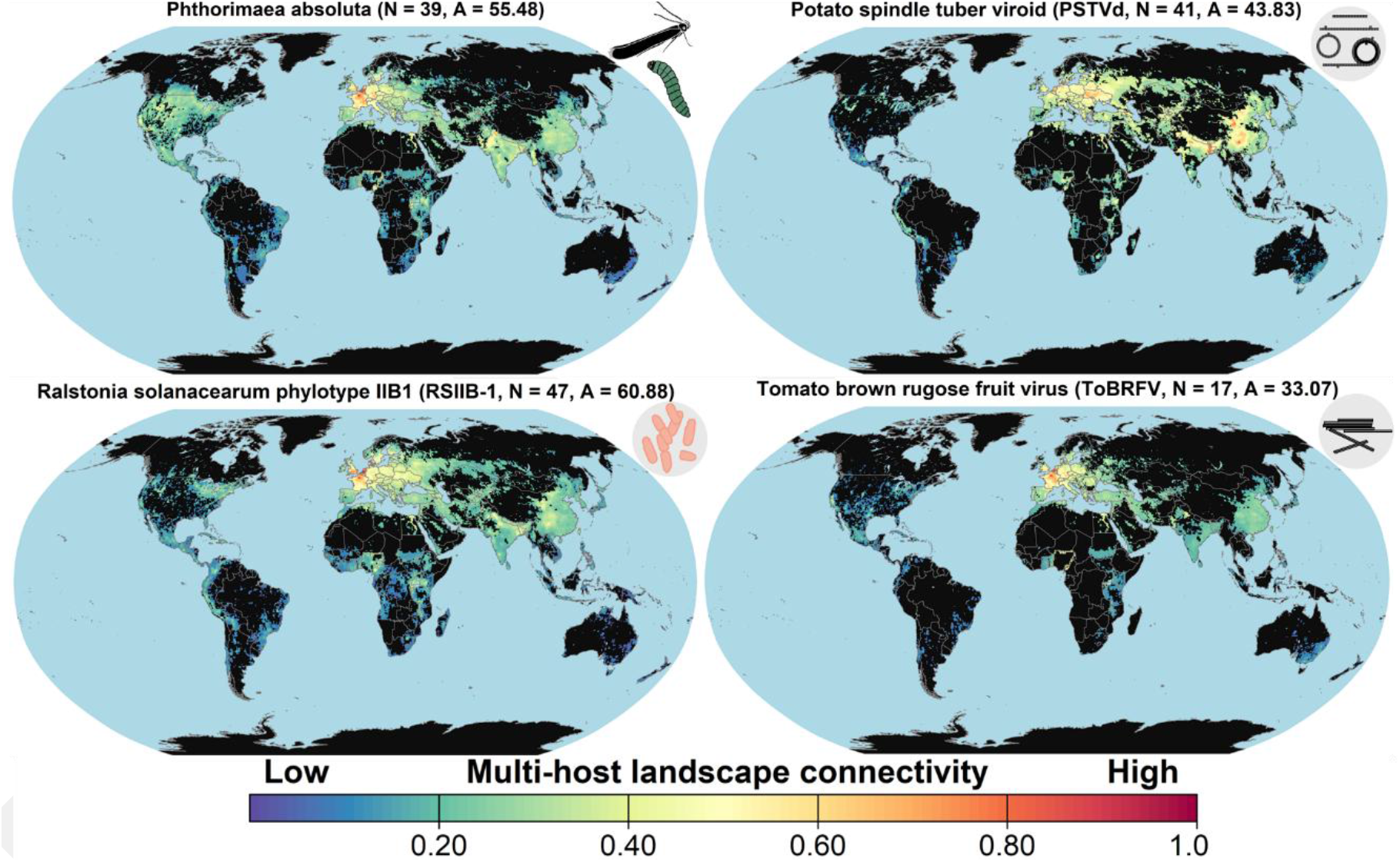
Global vulnerability hotspots for four invasive pests based on multi-host landscape connectivity. Color represents the magnitude of mean host landscape connectivity at each location, calculated across a range of likely dispersal parameters. In this global connectivity assessment, host community traits include the number of host species (cultivated, weedy, or wild species) for which natural infection has been demonstrated (N), and the land surface area with hosts available (A, in million square kilometers). Grid cells in black are locations where the host is not reported.

Host communities in the Americas and Africa are spatially fragmented for PSTVd, RSIIB-1, and ToBRFV. Despite this structural habitat fragmentation, the likelihood of pest movement due to functional host connectivity in the Americas and Africa is proportionally greater than if we consider only host availability. For example, California, Burundi, Rwanda, and a western area in Kenya have particularly high host connectivity for PSTVd, RSIIB-1, and *P. absoluta*. These highly connected host communities are potential pathways for spread if the pest reaches them.

In the Americas, there is a conterminous host belt through the Andes for the potential natural dissemination of each target pest. Panama is possibly a non-host disconnection for the natural spread of PSTVd, ToBRFV, and *P. absoluta*. In contrast, a contiguous host landscape in Central America is expected to act as a structural and functional bridge zone for the gradual spread of RSIIB-1 between North America and South America. Reported host availability is scattered throughout Africa, especially for PSTVd and ToBRFV. However, a landscape along the northern and eastern borders of Nigeria has high host connectivity for each target pest (Fig. 5). The reported host landscape fragmentation in Africa^25,26^ suggests the need for future assessments of unreported host distributions that could influence pest spread.

Considering only cropland substantially reduced the estimated invasion vulnerability compared to a multi-species host assessment for each target pest. The number of host species was weakly negatively or positively correlated with functional host connectivity (Spearsman’s *r*_*s*_ = –0.04, *p* = 1.486e-13 for PSTVd; *r*_*s*_ = 0.31, *p* < 2.2e-16 for RSIIB-1; *r*_*s*_ = 0.19, *p* < 2.2e-16 for ToBRFV; and *r*_*s*_ = 0.49, *p* < 2.2e-16 for *P. absoluta*). A Kolmogorov-Smirnov test indicated that multi-host connectivity tends to be higher in locations where the pest is reported than elsewhere (p < 0.001, Table S6), consistent with a likely role of multi-host connectivity in driving the spatial occurrence of each pest. These findings indicate that cross-species transmission is likely where crop ranges overlap with non-cultivated host species, a pattern that GIRAF is the first to quantify globally.

### A multi-dimensional biogeographic synthesis of pest invasions

Each component of GIRAF provides a different perspective on invasion risk, and decision-makers can use a single-factor map as a first approximation of a pest’s invasion potential when other information is lacking. Each map represents a snapshot of a pest’s potential geospatial distribution and a dimension of its ecological niche. The core innovation of GIRAF 1.0 is the integration of these individual perspectives into a Biotic-Abiotic-Migration (BAM) model^5,12^ (Fig. 6). By synthesizing trade-driven introduction pressure (migration niche), environmental suitability (abiotic niche), and host connectivity (trophic niche), this multidimensional analysis offers a holistic assessment of global invasion risks across these drivers. To our knowledge, GIRAF is the first global model for pest invasions that combines (a) pest-specific ecological niches, (b) real-world, spatially explicit data about major drivers of pest spread, and (c) explicit integration of natural and agricultural ecosystems.

**Fig. 6.**
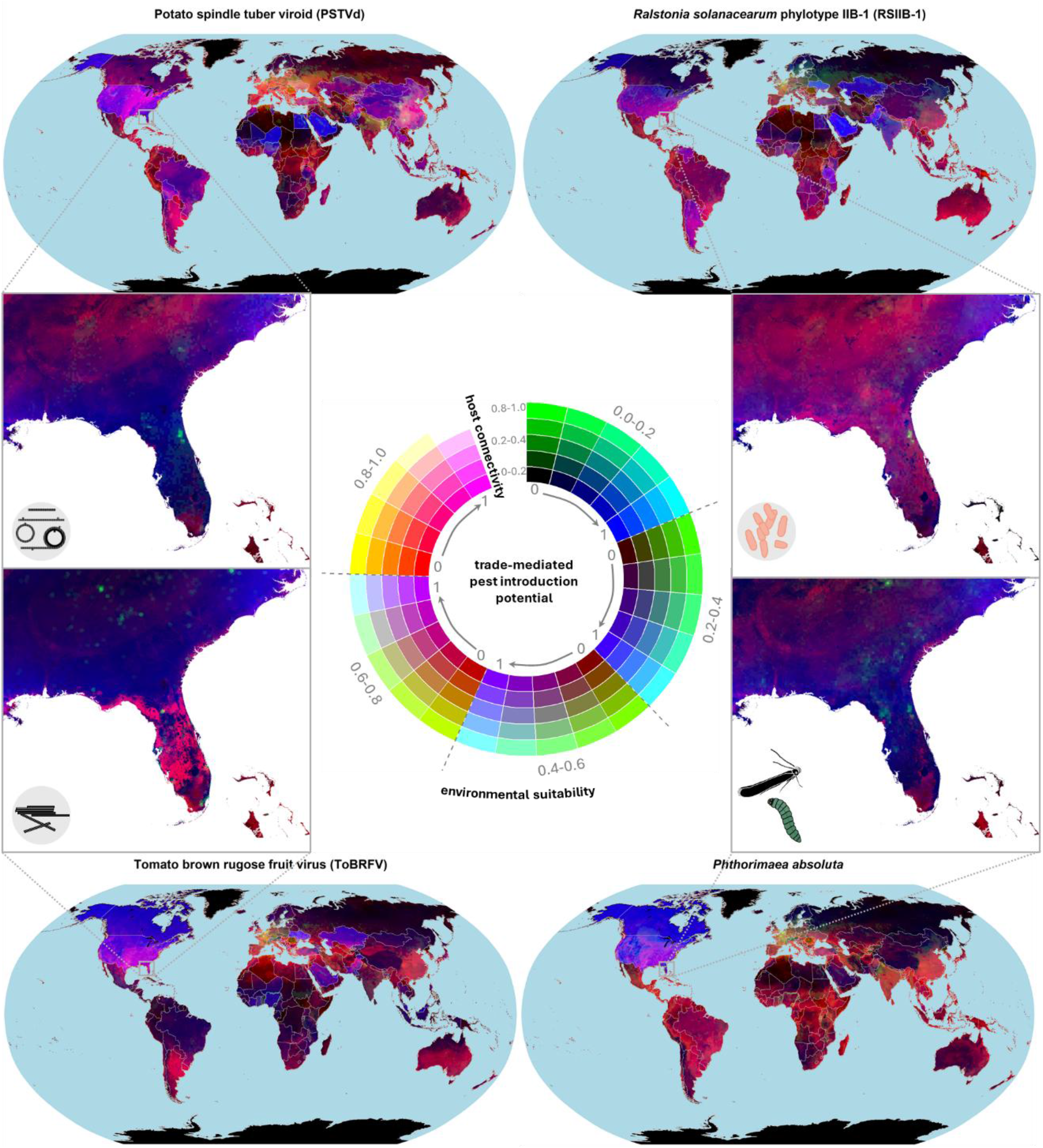
Worldwide pest invasion vulnerabilities based on a multi-dimensional assessment. In these multivariate choropleth maps, the intensity of each primary color represents the potential invasion level of a target pest based on environmental niche (red spectrum), host landscape connectivity (or trophic niche, green spectrum) and introduction potential via international trade of crop commodities (blue spectrum). Grid cells in black indicate areas with no invasion vulnerability (e.g., Antarctica), and pale-yellow grid cells indicate the highest vulnerable areas for a pest invasion where high levels of the three factors coincide. As an example, grid cells in orange have high environmental suitability but relatively low levels of host connectivity and trade-mediated introduction potential.

Our multidimensional analysis reveals the complexity of biological invasions: the highest levels of these three risk factors rarely coincide globally. We identify central Europe, India, and northern China as primary global epicenters where all three risk dimensions converge at high levels for the target species (Fig. 6). Conversely, in the USA, there is high introduction potential and environmental suitability, but host distribution may be a limiting factor for pest establishment.

### Multiscale lens for actionable biosecurity

GIRAF’s lens scales from global policy to local biosecurity, accounting for likely scale-dependent processes in biological invasions. At a finer spatial resolution (∼5 km), the framework identifies actionable priorities for local agencies. In the Southeastern USA, for instance, GIRAF reveals that northern Florida has moderate but non-negligible levels of all three risk factors for each target pest (Fig. 6). This granularity allows decision-making to focus either on large-scale warnings or spatially explicit surveillance, demonstrating that GIRAF provides a scalable, data-driven foundation for One Biosecurity^7^.

## Discussion

Protecting plants from invasive pests is key to safeguarding agroecosystem provisioning and natural ecosystem functions, especially in response to 21st-century global challenges like resource depletion, plant pandemics, and extreme weather events. GIRAF 1.0 identifies which areas are likely most vulnerable to pest invasions and most important to invasive spread, and thus candidate priorities for action. These atlases of invasion risk can serve as strategic tools when planning for global invasion preparedness, spatially explicit surveillance prioritization, and risk mitigation. By integrating four ecologically important drivers of the global spread of non-native pests, GIRAF 1.0 represents a key advance in invasion risk assessment.

Here, GIRAF identified potential vulnerability hotspots for four pests of global concern, combining new pest-specific biogeographic models of trade-mediated introduction potential, environmental suitability, and host landscape connectivity for a contemporary timespan. Our results provide the first quantitative assessment of invasion vulnerabilities for plant pests across these geographic factors globally. We provide new data-driven evidence that host communities occupy areas ranging from ∼33.1×10^6^ km^2^ for ToBRFV to 60.8×10^6^ km^2^ for RSIIB-1. In other words, ∼22 to 37% of 111×111 km cells on Earth’s land surface have some host available. Despite the implementation of phytosanitary policies, more than 115 countries remain highly vulnerable to (re)introductions, given the global trade of high-risk commodities associated with each pest. The USA ranks among the top five countries with trade-mediated (re)introduction vulnerability, and large areas of the country are environmentally suitable for pest invasions.

These findings will tend to have higher uncertainty about the invasion risk of these pests in regions where host availability is not reported, where climate variability, wind patterns, or human mobility are important factors, where informal trade in planting materials is common, and where international borders are present^14,27,28^. Despite these uncertainties, our findings provide inputs for planning spatially explicit proactive responses to and plant protection strategies against ongoing and new pest outbreaks (Fig. 1). We propose that responses should include: (1) proactive surveillance targeting regions where each pest is unreported, but which have high invasion vulnerability (Figs. 2-4); (2) making international trade networks safe through enhanced pest-specific biosecurity in over 115 countries (Figs. 2 and 3); and (3) regional management efforts to break high host connectivity, particularly where solanaceous crops geographically overlap with non-cultivated host species (Fig. 5-6).

Together, enhanced invasion and pandemic prevention, preparedness, and response are needed given the ongoing spread and impacts of these pests. GIRAF contributes to effort prioritization by predicting previously unquantified yet widespread invasion risks for each target pest. Long-term transnational strategies for (pro)active surveillance and risk mitigation can reinforce current nationally coordinated plant health systems^14,29^ in order to prevent future introduction, establishment, and local spread of these pests of global concern. Effective management of these transboundary pests also calls for collective support from the private sector (ornamental and food crop industries), conservation agencies, and NPPOs. This globally coordinated system for invasion mitigation needs to be adapted by biosecurity agencies and plant industries to regional circumstances.

GIRAF 1.0 has practical and cross-disciplinary relevance beyond the four globally concerning pests in this article. It can be applied to other target plant, animal, or microbial taxa, provided that sufficient data is available to reproduce the geographic synthesis. Widespread implementation of GIRAF 1.0 depends on readily available, interoperable pest information systems and timely financial support^29^. Global biosecurity will be strengthened if countries build and share local pest-specific data. GIRAF 1.0 integrates across multiple factors of pest spread, but does not yet fully incorporate the complex reality of biological invasions^30^, lacking pest species interactions with natural enemies, evidence from population genetics, inherent stochasticity, and fine-scale spatiotemporal dynamics. Invasion risk frameworks like GIRAF generally lack explicit quantitative impact assessments for multifaceted outcomes globally, such as crop yield losses, agriculture profit reduction, plant biodiversity losses, or environmental impacts^2^. These research frontiers for pest information systems challenge any invasion framework designed to inform timely interventions for real-time geographic monitoring prioritization. GIRAF 2.0 should tackle these grand challenges in invasion science as biogeographic pest information systems improve. We hope GIRAF 1.0 serves the scientific community as a reference model to design future global risk assessments for the thousands of potential invasive species currently threatening Earth’s plant ecosystems.

## Methods

GIRAF incorporates four main steps for quantifying invasion risk: (i) defining ecologically or epidemiologically important risk factors, (ii) collecting or compiling fine- or broad-scale data related to these risk factors, such as dispersal pathways, geographic occurrence of species, and host range, (iii) selecting and (re)training the biogeographic model(s) based on digitally accessible information, and (iv) generating evidence-based maps of candidate priorities for surveillance and mitigation. While GIRAF can serve as a general framework for future assessments, our choice of specific biogeographic models to quantify environmental suitability, host connectivity, and introduction risk is not prescriptive. Depending on the goals of risk assessments, other biogeographical models can be used to quantify invasion risk for a pest species.

When implementing GIRAF, risk analysts, policymakers, and biosecurity practitioners can provide periodic feedback on each component and iteratively fine-tune the resulting spatial projections of pest invasion vulnerability, particularly if relevant information like informal trade of agricultural commodities is only privately or unofficially documented. As for all models, the quality of outputs is only as good as the quality of the data input. No single database used here is bias-free. We focus on the four plant pests as real-world case studies because they are invasive species priorities for the US and globally; selection of pest and disease priorities is an expert-driven process external to GIRAF^11^[Supplementary Text 1].

### Data assembly for ecologically important species traits

We built (i) a geographic distribution dataset including the reported countrywide extent of each pest species, the earliest year of the pest collection or detection in the country, the first year of the country report publication, and georeferenced presence records, wherever available (Data S2-7); and (ii) a host-parasite association list including plant species naturally or experimentally infected by the pathogen or infested by the pest, and the reported countries of these associations (Data S8). These datasets present a comprehensive compilation of data from publicly available reports through 2023, including journal articles and official reports from NPPOs. Despite this extensive data compilation, global systematic sampling or highly standardized reporting for these pest species are rare. While the spread of pests at large spatiotemporal scales cannot always be systematically represented by small-scale field and laboratory experiments^31^, ‘national- or continental-scale controlled trials’ are generally not realistic^32^. Despite their systematic incompleteness, geographic bias, and often convenience-based sampling, observational distribution data serve as a primary source of empirical information for mapping the potential spread of invasive pests across broad-scale crop-growing regions.

For each natural host species listed in the host-parasite association spreadsheet, but unavailable in the CROPGRIDS dataset^26^, we created maps of relative host density. We manually downloaded species-specific georeferenced occurrence records from the Global Biodiversity Information Facility (GBIF) database on July 5^th^, 2024^25,33,34^. On a global map with grid cells of approximately 2.3 km at the equator (1.25-minute spatial resolution), we assigned each grid cell the square root of the number of presence records for host species, or 0 if there were no georeferenced records. These global maps represent the geographic distribution of individual plant species at relatively high spatial resolution and are expected to be highly biased in areas with lower sampling effort. We thus aggregated each map at ∼55 km resolution (0.5° per grid cell), calculating the mean grid cell values at coarse resolution, to partially reduce sampling bias^33^. These maps represent a first approximation of relative host density; future approaches could train species distribution models to produce more accurate maps for each host. For cultivated host species, we obtained global maps of crop-specific harvested areas from the CROPGRIDS dataset, which incorporates production statistics that are not included in GBIF records of crop species.

#### Spatial coverage

Each analysis targeted two geographic extents. Global analyses are presented at a spatial resolution of 0.5 °. Each trained model also produced focused risk maps for each pest, for Florida, Alabama, Georgia, and South Carolina, resampled at 3-minute spatial resolution.

#### Mapping invasion vulnerability based on species bioclimatic modeling

We obtained global maps of the 19 bioclimatic variables from CHELSA Bioclim, representing climate conditions for 1989-2013^35^, and of 12 soil properties from SoilGrids 2.0, representing edaphic conditions at 15-30 cm standard depth^36^. We also gathered 13 available maps of physical accessibility of land areas worldwide^37^. Four accessibility maps indicate travel time to airports and maritime ports, each representing one of four port sizes. We assigned individual weights to each port size map because each may have a different degree of importance for the entry of commodities and associated pests (Table S2). We then built an overall accessibility index to ports (*A*_*p*_) as a weighted average of accessibility to individual port sizes. The remaining nine maps characterize travel time to urban and rural locations, each aggregated at one of nine settlement classes (population size classes). We assigned a different weight to each settlement class to calculate a weighted average of overall access to cities (*A*_*c*_) across the nine settlement classes (Table S3).

We trained four probabilistic machine learning algorithms – MaxEnt, random forest, XGboost, and logistic regression –commonly used for presence-only data ^38,39^. Each algorithm represented a correlative species distribution model (SDM), in which the response variable was 1 for reported georeferenced presence records and 0 for pseudo-absences (i.e., background points randomly selected from the world land area). Each SDM was initially trained and evaluated on the following selected predictor variables: annual mean temperature, mean diurnal range in temperature, isothermality, annual precipitation, precipitation seasonality, chemical soil properties (pH, and soil organic carbon content), physical soil properties (clay, sand and silt content), port accessibility (*A*_*p*_) and city accessibility (*A*_*c*_). These predictors are a subset of all variables available in each dataset, allowing us to avoid multi-collinearity, while still maintaining substantial variation in ecologically relevant covariates of invasive species distribution. These initially trained SDMs indicated *A*_*p*_ as the most important variable explaining the reported distribution of each invasive species (54%, 27%, 72%, 53% contribution in presence predictions for PSTVd, RSIIB-1, ToBRFV, and *P. absoluta*, respectively, based on MaxEnt; Data S1). However, we excluded the contribution of accessibility in the final predictions by each SDM to provide a more explainable, deterministic approach. In this approach, we analyzed both *A*_*p*_ and *A*_*c*_ along with introduction vulnerability due to international trade to explicitly consider the individual ecologies of each invasive pest (see below).

This multi-model approach was used to generate a global map of ensemble predictions, equally weighting the spatial projections of these four SDMs, as a quantitative consensus approximation of abiotic environmental suitability for each invasive species. These species-presence predictions based on occurrence-environment associations are an initial and provisional approximation to environmental suitability for a species. True mechanistic ecological interactions between abiotic environmental conditions and occurrence data of these invasive pests have generally not been characterized quantitatively. Importantly, some locations are likely to have a higher climate suitability than predicted by the ensemble approximation, as will likely be discovered as each pest continues invading new environments and geographical spaces. We lack a quantitative understanding of how edaphic or climate conditions directly restrict or facilitate geographic occurrence. Climate effects on each stage in the life cycle are not currently available for most pest species, particularly for plant viruses that are not transmitted by vectors, such as ToBRFV. However, once this ecological information becomes available, process-based, component-based, or mechanistic models for these pest species can be preferentially used. These models would explicitly incorporate direct climate effects on pathogen distribution or a species’ physiological response to environmental conditions.

We adjusted our probabilistic bioclimatic ensemble with known environmental parameters for each species. We used Shelford’s law of tolerance to adjust ensemble predictions of environmental suitability for *Phthorimaea absoluta* and RSIIB-1. The law of tolerance states that an organism’s success is determined by a set of minimum, optimum, and maximum environmental conditions^40^. Using this ecological principle, we applied a generalized beta distribution model to project the potential invasion risk as a response function dependent on temperature^41^. In this thermal niche model, invasion risk (*r*(*T*)) depends on three cardinal temperatures for a species’ population development (Data S12): the minimum temperature (*T*_*min*_), optimum temperature (*T*_*opt*_), and maximum temperature (*T*_*max*_). We used the monthly mean temperature of each location (*T*_*x,y,m*_, where x and y are the geographic coordinates and *m* is the month considered) to estimate pest invasion risk locally:

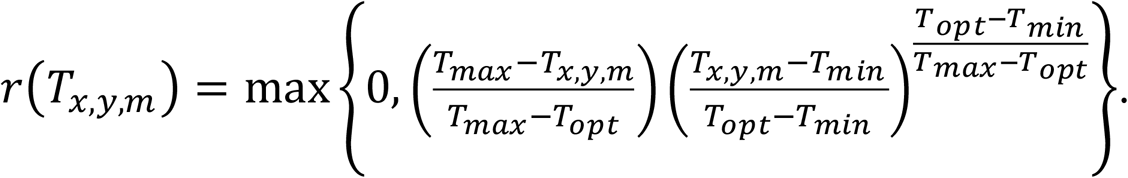

Invasion risk is highest at locations where *T*_*x,y,m*_ = *T*_*opt*_, decreases at temperatures higher or lower than *T*_*opt*_, and reaches zero beyond critical thermal limits tolerated by a species (below *T*_*min*_ or above *T*_*max*_). This temperature-driven physiological response is common in arthropods, plants, nematodes, fungi, and bacteria, and applies to *P. absoluta* as well as the cold-tolerant RSIIB-1 strains^42-44^. Here, cumulative pest invasion risk in a location over a year *r*(*T*_*x,y*_) is proportional to the sum of *r*(*T*_*x,y,m*_) for each month. We regarded climatically unfavorable locations as those with *T*_*x,y,m*_ < *T*_*min*_ or *T*_*x,y,m*_ > *T*_*max*_, defining geographically possible thermal range frontiers of a species.

Surface water such as rivers may serve as an aquatic habitat for survival, evolution, and dissemination of plant pathogens ^45-47^. We incorporated river networks in GIRAF as a possible plant health risk and a dispersal pathway for RSIIB1 (Supplementary Text 1). Using the HydroATLAS database ^48^, we calculated the mean river water discharge as a relative proxy for the likelihood that RSIIB-1 would disperse to any climatically suitable location globally.

No information was available about the direct effect of environmental variables on disease caused by ToBRFV and PSTVd. Here, environmental risk modeling for these species is based solely on the machine-learning ensemble.

### Mapping (re)introduction vulnerability based on international trade of crop commodities

As a key component for developing safe trade strategies, we characterized the structure of trade networks to identify potential geographic paths of pest spread and the relative vulnerabilities of locations to potential (re)introduction(s) for each pest species^8,9,29^. Hereafter, we define invasion risk as the likelihood that a pest or pathogen (i.e., hazard) potentially reaches or occurs in a host location^10,12,21^. We use the term invasion vulnerability to make clear that our analysis focuses on the risk of pest or pathogen entry into a country, rather than the risk that a country might pose as a source of inoculum through trade. In all our analyses, we used relative indices to estimate the likelihood of spread of an invasive species as approximations for invasion risk. In the global trade networks, specifically, we quantify the relative likelihood of potential spread of an invasive species through the international trade of agricultural commodities.

We gathered information about the trade volume of crop-specific commodities between each pair of countries, based on bilateral import reports in the World Trade Organization (WTO, https://stats.wto.org/) dataset for 2005-2019 and Volza (https://www.volza.com/) dataset for 2023. Our proxy for host availability within a country was the harvested area of crop species reported to be natural major hosts of each pest (Data S8), for crop species available in the FAOSTAT dataset^23^. To account for the potential effect of pest-associated trade policy landscapes, we also obtained information on international biosecurity measures targeting specific pest species, whenever available. We compiled information on the geographic extent of each pest within a country (Data S2-7), based on available reports in CABI Compendium (https://www.cabidigitallibrary.org/journal/cabicompendium), EPPO Global Database (https://gd.eppo.int/), and extensive literature review. In these international trade networks, nodes represent countries, and link weights indicate the relative potential of pest spread between countries.

We propose a trade-mediated introduction index (or *P*(*τ*_*i*→*j*_)) for potential accidental pest movement from an exporting country *i* to an importing country *j* as a quantitative proxy characterizing pest invasion risk in trade networks. For any pair of trading countries, *τ*_*i*→*j*_ combines explicitly and quantitatively the geographic extent of a pest species in each of the trading countries, the host availability in each of the trading countries, the trade volume of crop-specific commodities between the countries, and, when available, pest-specific biosecurity measures implemented by trading countries. Supplementary Text 2 provides details of the methodological approach, mathematical formulations, theoretical assumptions, and available datasets used to estimate *P*(*τ*_*i*→*j*_) through international trade networks. We modeled the joint relative likelihood that none of the exporting countries would introduce the pest species into a target importing country *j* as 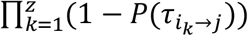, where *z* is the number of countries exporting a crop-specific commodity to country *j*. Finally, we assumed that the joint risk (*I*_*j*_) that the target pest is introduced into a country from any exporting countries is directly proportional to 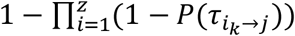. As an alternative measure, we also calculated three network metrics to characterize the potential introduction risk of a pest species to a country (i.e., *I*_*j*_): node in-strength, betweenness centrality, and eigenvector centrality. These network metrics have been important in predicting pest or pathogen transmission in epidemic networks^49-51^.

Our geographic risk analysis of the potential (re)introduction of the four invasive species focused on individual networks of the reported international trade of specific agricultural commodities. For PSTVd, we analyzed networks of international trade of seed potato (i.e., potato tubers for 2005-2019) and planting material of *Brugmansia* (2023). For RSIIB-1, we evaluated the international trade of seed potato, tomato fruit, pepper fruit (2005-2019), and geranium planting material (2023). For ToBRFV, we built individual networks of international trade of tomato fruit (2005-2019), tomato seed, and pepper seed (2023). For *Phthorimaea absoluta*, we assessed the international trade network of tomato fruit (2005-2019). These target commodities are important for the international dispersal of the pests of interest, given their reported specific association with the interception of these pest species (Data S10). We also focused the (re)introduction risk analysis on the international trade of these fresh crop commodities because of their potential higher likelihood in the geographic diffusion of these invasive pests, excluding processed agricultural products which may have a negligible risk. Future risk analyses could include explicit information on other primary dispersal pathways of these invasives in the longer term. Information about the international trade of crop-specific seed or planting material over multiple years is not available publicly. To include the potential effects of implemented policies in our assessment, we modeled a reduction in the introduction risk of 10% from countries with market access to the United States that imposed import biosecurity requirements. The list of countries with pest-specific biosecurity regulations is available in the 2024 Federal Order for U.S. imports of tomato and pepper seeds for ToBRFV, and the 2023 Federal Order for U.S. imports of tomato fruit for *P. absoluta* (Data S3). We also included an analysis focused on the vulnerability of the USA and countries in the Caribbean region to introduction of each pest species associated with commodity imports, providing a regional assessment.

We used country-level interception data for each pest species as a test dataset to validate the (re)introduction vulnerability analysis (Data S10). To build the introduction model above, we used this data to determine which agricultural commodities are likely important for the international spread of each pest (Table S5). Below, we used pest interceptions in specific countries that were not used in the model construction, except when interception are the only information available for the status of a pest. We assessed the precision of the (re)introduction vulnerability analysis, that is, the ratio between (a) the number of countries where the pest has been intercepted on imported agricultural commodities and introduction vulnerability was non-zero (true positives) and (b) the number of countries where the pest has been intercepted on imported agricultural commodities. Our analysis had a precision of 1, including all countries where the pest has been intercepted (Table S7). Using a Kolmogorov-Smirnov test, we also evaluated whether countries where the pest has been intercepted have a higher (re)introduction vulnerability than any other countries. The KS test did not detect evidence for higher or lower introduction vulnerability in countries where the pest has been intercepted (Table S7). There was not evidence to reject the null hypothesis (***p*** > 0.27 for all pests), so a lack of evidence that higher vulnerability values were associated with pest (re)introduction.

### Mapping invasion risk based on accessibility to ports and cities

Ports likely play a pivotal role in the (re)introduction of plant pests to a region, as they may serve as primary entry points of pest-associated agricultural commodities^3,9^. Geographic proximity to ports generally increases the risk of introducing non-native arthropods and pathogens^3,52^ and our SDMs indicated a potential major role of access to ports in the geographic distribution of each target pest. Thus, our models incorporate accessibility of croplands to ports or urban areas in a region as increasing (re)introduction vulnerability to plant pests. Likewise, accessibility of croplands to cities, in general, may increase invasion risk associated with urban agricultural landscapes and the final destination of commodities^53^. Specifically, we treated invasion risk associated with accessibility to ports and cities as occurring in a pattern analogous to species richness resulting from species-accumulation models, where the cumulative number of species scales in an exponential pattern with sample size, area, or intensity^54^. We modeled invasion risk of cropland locations in a region as increasing nonlinearly with accessibility to ports (*A*_*p*_) as *I*_*p*_ ∝ 1 − exp (−1/*log*(*A*_*p*_)) or with accessibility to cities (*A*_*p*_) as *I*_*c*_ ∝ 1 − exp(−1/*log*(*A*_*c*_)), where *I*_*p*_ ∈ [0,1] and *I*_*c*_ ∈ [0,1].

For each invasive species, we generated a map integrating the vulnerability of each country to a pest’s introduction through international trade and based on the accessibility of croplands to ports (*I*_*j*_ × *I*_*p*_). This resulting map aims to disaggregate the accidental introduction potential of pest species at the national level (*I*_*p*_) into likely domestic distribution of imported agricultural commodities and their associated pests across initial entry locations. These maps of invasion risk can be fine-tuned in future geographically explicit evaluations, as domestic distribution of commodity trade and local spread of associated pests may vary geographically among commodity types^52^. Information about origin location, ports of entry, and final city destination of imported agricultural commodities specifically associated with a target pest species is not publicly available, to our knowledge.

### Mapping invasion risk based on multi-host landscape connectivity

Our target invasive pests have multi-species host ranges. Their potential distribution in regions where a host plant is unavailable is constrained^30^, but the risk of pest spread tends to be higher where susceptible host populations are homogeneously and densely distributed across the landscape^55-57^. Geographic host distribution is a critical risk factor when accounting for biotic interactions more realistically in ecological niche modeling of plant disease^55-57^. We categorized each plant species reported to be naturally infested by a pest as a primary or secondary host (Data S8; Table S4). Natural secondary host species were included as playing at least a minor role in pest persistence or survival. To map the geographic distribution of multiple natural host species, we used a stacked host distribution modeling approach^12^, summing the relative density of major and secondary host(s) to produce a global map of cumulative host density for each invasive pest. In these stacked host maps, we considered the potential minor role of secondary host species in pest invasion risk and potential spread by multiplying their host densities by a tenth^9^. For crop species being host of a pest species, our analysis included only locations represented by 3-minute cells having ≥ 4 hectares of cropland (a host density threshold of ∼0.1%), incorporating a rare-species advantage against density-dependent diseases in excluded host locations^55^. We then aggregated these resulting maps to a coarser spatial resolution so that each grid cell represented ∼55.5 km × 55.5 km at the equator.

Using these global maps of cumulative host density as inputs in the geohabnet R package version 2.2^58^, we evaluated the host landscape connectivity for each pest species^21^. geohabnet is a component of the R2M Plant Health Toolbox for rapid risk assessment to support mitigation of pathogens and pests (www.garrettlab.com/r2m). Using geohabnet, we estimated the relative likelihood of pest movement (*σ*) between host locations *i* and *j* using two generic gravity models for species dispersal^59-61^: 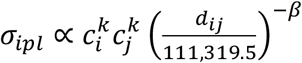 for the inverse power-law model and 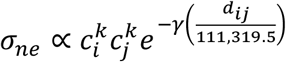 for the negative exponential model. In these global dispersal models, potential pest movement (*σ*_*ipl*_ or *σ*_*ne*_) depends not only on the product of relative abundances of susceptible host species in both locations 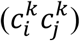, but also on the probability of a pest moving between host populations given their physical distance (*d*_*ij*_). We set 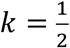 to account for potential nonlinear associations between host density and pest invasion risk^62^. We compiled *β* and *γ* dispersal parameter values that were empirically estimated for a diverse set of plant pathogens and arthropod pests (Data S11). We used this dataset to calculate the first quartile, mean, median, and third quartile of each dispersal parameter across pest species, representing a parameter space of likely pest spread scenarios. We evaluated these typical parameter values *β* = (0.9, 1.5, 1.7, 2.1) and *γ* = (0.02, 0.08, 0.36, 0.24) in a sensitivity analysis to account for uncertainty about pest movement. Species-specific dispersal parameter values are unavailable for these target pests (as well as for most pest species). We then built pest invasion networks, where a node represented a host location, and link weights indicated potential pest movement between host locations (*σ*_*ipl*_ or *σ*_*ne*_).

We calculated host landscape connectivity using six standard network metrics in epidemiology and invasion ecology^18,49,63-66^: betweenness centrality, closeness centrality, eigenvector centrality, node strength, PageRank centrality, and the sum of nearest neighbors’ node degrees. Here, global maps of invasion risk represent the multi-host landscape connectivity for a target invasive pest, averaged across two gravity models, eight typical dispersal parameter values, and six standard network metrics, each weighted equally. Host landscape connectivity quantifies the relative likelihood that an invasive pest, if it reaches the target host location, will spread to all its direct and indirect neighbors in the network.

In a separate analysis, we treated pest survival as higher in areas with higher host species richness than in areas with only one host species^12^. We used georeferenced occurrence data for each pest species as an independent dataset to validate the multi-host connectivity estimates. We used two metrics to assess model performance based on presence-only data. First, precision is the ratio of the number of grid cells in which the target pest is reported present and multi-host connectivity is nonzero (true positives) to the total number of grid cells in which the target is reported present (true positives + false negatives). Second, we assessed whether multi-host connectivity is higher in locations where the pest is reported to be present than elsewhere. To assess this hypothesis, we conducted an asymptotic two-sample Kolmogorov-Smirnov test. The multi-host connectivity analyses had good precision (from 0.68 to 0.87; Table S6).

## Acknowledgments

The opinions expressed in this article are those of the authors and do not necessarily reflect the views of the USDA. We appreciate insightful comments from Romaric A. Mouafo-Tchinda, Ashish Adhikari, Manoj Choudhury, Jacobo Robledo, and Mathews Paret.

## Funding

USDA Animal and Plant Health Inspection Service (APHIS) Cooperative Agreement AP21PPQS&T00C195

USDA Animal and Plant Health Inspection Service (APHIS) Cooperative Agreement AP22PPQS&T00C133

USDA Animal and Plant Health Inspection Service (APHIS) Cooperative Agreement AP24PPQS&T00C076

USDA National Institute of Food and Agriculture (NIFA) grant 2022-51181-38242 USDA National Institute of Food and Agriculture (NIFA) grant 2024-67013-42781 USDA Hatch grant 1023861

## Author contributions

Conceived and designed the experiments: AIPS, GC, KS, ES, PS, KAG

Performed the experiments: AIPS

Analyzed the data: AIPS, OB, GC, NSD, BAE, AH, TMLP, JDM, CP, KS, ES, PS, YT, HT, YW, KAG

Contributed materials/analysis tools: AIPS, TMLP, OB, KAG

Wrote the paper: AIPS, OB, GC, NSD, BAE, AH, TMLP, JDM, CP, KS, ES, PS, YT, HT, YW, KAG

## Data and materials availability

A template for processing R scripts for each analysis in this study is publicly available on GitHub: https://github.com/AaronPlex/pestradenet for international trade networks, https://github.com/AaronPlex/pest-env-sdm for environment-based species distribution models (SDMs), and https://github.com/AaronPlex/multi-host-nets for host connectivity analysis. All data are available in the main text or the supplementary materials. All unpublished datasets supporting the results and reproducibility of this study are available as supplementary data. All published datasets used in this study are correspondingly cited.

## Supplementary Materials

This manuscript includes the following Supplementary Materials: Supplementary Text 1 to 2, Figs. S1 to S5, Table S1 to S2, and Data S1 to S10.

