## Supplemental Material for "GIRAF 1.0: A unified global framework to anticipate plant pest invasions"

Aaron I. Plex Sulá *et al.*

**This PDF file includes:**

Supplementary Text
Figs. S1 to S5
Tables S1 to S2
Data S1 to S10

**Supplementary Text 1**

i) The national and regional context of pest surveillance and mitigation systems

Pest invasion preparedness and risk mitigation involve a series of plant biosecurity activities [1-3]. Pre-invasion biosecurity activities require periodic inspection, interception, and surveillance of potential pest entry points and likely source regions. However, complete exclusion of invasive pests in new geographic areas is often as challenging as "finding a needle in a haystack" [4]. Implementing pre- and on-border biosecurity activities is economically and logistically feasible in only a small fraction of imported commodities and potential pest arrival locations in a host landscape [5-7]. Designing post-border surveillance is also critical for early and rapid response to plant pest invasions regionally. For example, strengthening biosecurity links between Florida and the Caribbean region is particularly important because of their relatively similar climate, high production capacity, high intraregional trade, and geographic proximity.

In the conterminous United States, USDA's Animal and Plant Health Inspection Service's Plant Protection and Quarantine (APHIS PPQ) and the Department of Homeland Security's Customs and Border Protections (DHS CBP) are federal agencies in charge of conducting, developing and implementing plant biosecurity activities, from offshore detection to local management of exotic pest invasions [1]. Likewise, the Cooperative Agricultural Pest Survey (CAPS) program conducts non-native plant pest surveys to protect American agriculture and natural resources (https://www.aphis.usda.gov/plant-pests-diseases/caps). CAPS is a national network of state and federal cooperators and stakeholders that focuses on early detection to prevent establishment of exotic plant pests [8]. There are also statewide agencies that safeguard agriculture and natural resources against local plant pest invasion and initial epidemics, such as the Florida Department of Agriculture and Consumer Services (FDACS). With an annual budget of ~US$55 million, FDACS Division of Plant Industry maintains phytosanitary programs both independently and in collaboration with the USDA, CBP and other entities, including the Cooperative Agricultural Pest Survey (CAPS) and plant inspections at agricultural interdiction stations along interstate corridors. APHIS invested approximately US$400 million in prevention of and preparedness for plant pest invasions annually [9]. Beginning with pest prioritization, a cross-sectoral committee organized by the National Plant Disease Recovery System (NPDRS) annually determined a high-priority list of invasive pests in the United States that need nationwide attention and investment [10]. Similarly, during a forum in collaboration with APHIS PPQ through the Greater Caribbean Safeguarding Initiative (GCSI), the Caribbean Plant Health Directors determined that *P. absoluta* and RSIIB are pests of concern to safeguard the greater Caribbean (https://www.cphdforum.org/). We focus on the four plant pests as real-world case studies because they are invasive species priorities in the US (https://approvedmethods.ceris.purdue.edu/), they have the potential to be introduced or have already been introduced to the United States and the Caribbean region, and they seriously threaten major solanaceous crops globally and locally.

ii) Understanding ecological or epidemiological traits of the targeted invasive pests

*Phthorimaea absoluta* can infest at least 42 plant species naturally (Data S1), with tomato being the major host species [11,12], and occasionally feeds on other cultivated solanaceous plants like potato, tobacco, pepper, and eggplant (Garzia 2009). *P. absoluta* is native to South America with an earliest known record in Peru in 1917, first reported in Spain in 2006 representing the earliest invasion of this species in Europe [11], and detected in Haiti in 2018, reaching the Caribbean Basin [13]. By 2022, *P. absoluta* had a global distribution with reported occurrence in 108 countries (Fig. S1; Data S1). The long-distance rapid spread of *P. absoluta* is unintentionally mediated by international trade of infested agricultural commodities, such as tomato fruit, in packaging materials, and in sorting facilities. Active dispersal is by short-distance flying, facilitated by wind currents [11].

*Ralstonia solanacearum* is a species complex clustered into four phylotypes, causing infectious disease in a wide range of economically important crops and non-cultivated plant species [14,15]. RSIIB strains are known to naturally infect 54 plant species and are the most frequently reported strains in this species complex causing bacterial wilt in potato, tomato, and pepper [16,17]. Non-cultivated plant species can serve as secondary or occasional hosts of RSIIB strains (Data S1). The RSIIB phylotype has a pandemic distribution [16], with occurrence reports in 62 countries (Fig. S1; Data S1), and a possible region of diversity in South America [18]. RSIIB was first detected in Florida in 2001 [19]. The dissemination of RSIIB strains over long distances occurs primarily and unintentionally through the international trade of infected ornamental plants like asymptomatic geranium cuttings, potato tubers, and tomato propagative material [17,19-21]. Surface water, such as contaminated rivers, ponds, and agricultural drainage, plays a key role in the survival of RSIIB strains when used for irrigation [21-24].

Tomato brown rugose fruit virus (ToBRFV) is an emerging transboundary pathogen in tomatoes, which was first discovered in Israel in 2014 [25] and Jordan in 2015 [26]. By 2024, there were reported outbreaks of ToBRFV in 50 countries (Data S1), posing a pandemic threat to the global tomato and pepper industries [27]. ToBRFV was detected in a tomato greenhouse in California in 2018 with eradication efforts implemented [28] and, more recently, was reported on imported tomatoes in grocery stores and community gardens in Florida [29-31]. Tomato and pepper are the main hosts of ToBRFV [27,32]. Yet, this virus can infect at least 17 solanaceous species naturally and 41 additional plant species in experimental inoculations (Data S1). Globally, ToBRFV has been intercepted at least 109 times in tomato and pepper seeds, tomato and pepper transplants, and tomato fruit (Fig. S1). These commodities are the main pathways of rapid international spread of this virus [32-34]. Locally, ToBRFV is mechanically transmitted via plant-to-plant contact, contaminated farming tools, workers' hands, bumblebees carrying the primary virus inoculum passively, and possibly irrigation water and contaminated soil from previous growing cycle [27,32,35].

*Potato spindle tuber viroid* (PSTVd) has a widespread but scattered global distribution with reported incursions in 50 countries and multiple interceptions of the pathogen in new regions over the past two decades (Fig. S1-2; Data S1). PSTVd is declared eradicated only in Canada and the United States, where the pathogen was first discovered in the 1920s [36]. PSTVd causes seed degeneration in potato tubers and occasionally rasta disease in tomatoes [37]. It naturally infects at least 51 cultivated and non-cultivated, mostly symptomless, plant species (Data S1). PSTVd has additionally 132 host species in inoculation experiments. PSTVd has been accidentally introduced into at least nine countries because of the international exchange of infected potato tubers, pepper or tomato seeds, and planting material of asymptomatically infected solanaceous ornamentals (Fig. S2).

**Supplementary Text 2**

i) Estimating pest invasion potential through global trade networks

We developed a mathematical model to characterize the potential movement of pathogen or pest species between countries, which explicitly accounts for six geographic risk factors: country-level host availability, within-country pest extent, international trade of agricultural commodities, proximity to ports and cities, national biosecurity capacities and pest-specific climate suitability. This analysis can be reproduced using the R code available at the following GitHub repository https://github.com/AaronPlex/pestradenet

Before describing how we integrated each risk factor in a quantitative way, it is important to mention the phenomenon of **directionality** in trade networks. Because agricultural trade activities are a directional process, pest movement through trade networks depends on the mode of trade activity, whether a country is exporting, re-exporting, importing, or exporting and importing agricultural commodities. Distinguishing which countries are sources, intermediaries ('re-exporters'), and destinations of agricultural commodities in a trade network is thus important to understand the potential direction of unintentional pest spread mediated by trade activities [e.g., [38-40]]. While importation of agricultural commodities is associated with the possible risk of pest introduction or re-introduction to a country, patterns in exportation of commodities help us understand the possible risk of pest release from a country. We also define **invasion potential** simply as the potential movement of a species from one location to another (a country represents a location in the case of international trade networks). For simplicity, this definition of invasion risk does not explicitly distinguish whether a pathogen/pest species is native or invasive in a country. Determining invasiveness of a pathogen/pest species is challenging and often unavailable for many plant pests and pathogens [41]. We also used the definition of **pest** from the International Standards for Phytosanitary Measure5 (ISPM 5) from the International Plant Protection Convention (IPPC). In accordance with this definition, we include arthropods and pathogens with a known negative impact on crops in our definition of plant pests.

ii) Invasion potential as a function of pest extent

The status of pest species within a country is usually expressed as nominal categories, such as those reported by the Centre for Agriculture and Bioscience International (CABI), European and Mediterranean Plant Protection Organization (EPPO), PlantwisePlus Knowledge Bank, and scientific journal articles. These categories are based on the reported presence or geographic range of a pest species, as well as the frequency of its occurrence within a country. A standardized metric that measures the full area occupied by a pest species within a country [42] is often not available in the literature, making it difficult to estimate how widespread a pest is within a country. We proposed an ordinal and relative ranking as a first approximation for quantifying the pest status or extent within a country (Table S1). This quantitative ranking results from the interpretation of each nominal category.

**Table S1. Proposed ranking for the geographic extent/status of pest species in a country.**

| Reported pest presence | Reported pest range or extent | Assigned score (Ɛ) |
|---|---|---|
| Present | Widespread (CABI, EPPO) | 5 |
| Present | Native | 5 |
| Present | Without details | 4 |

| | | |
|---|---|---|
| Present | Localized (CABI)<br>Restricted distribution (EPPO) | 3 |
| Present | Sporadic or episodic | 3 |
| Present | Exporting interceptions | |
| Present | Few occurrences | 2 |
| Present | Transient under eradication | 1 |
| Present | Transient under surveillance | 1 |
| Present | Transient nonactionable | 1 |
| Absent | Eradicated | 0 |
| Absent | Intercepted only | 0 |
| Absent | Formerly present | 0 |
| Absent | Never occurred | 0 |
| Absent | Simply not reported | 0 |
| Absent | Confirmed by survey, invalid record, intercepted only, pest eradicated, unreliable record (EPPO) | 0 |
| Not reported | Simply no reports are available | 0 |

We formulated an invasion potential index that depends on an expected relationship between the pest extent in the exporting or source country ($\varepsilon_i$) and the pest extent in the importing or destination country ($\varepsilon_j$). This expected relationship is given by $\frac{\varepsilon_i}{\varepsilon_j+1}$, so that the risk of a pest moving between any pair of countries based on their pest extents ranges between 0 and 5 (see the heatmap below). In this heatmap, dark red cells represent a maximum invasion potential (5), and dark blue cells represent a minimum invasion potential (0).

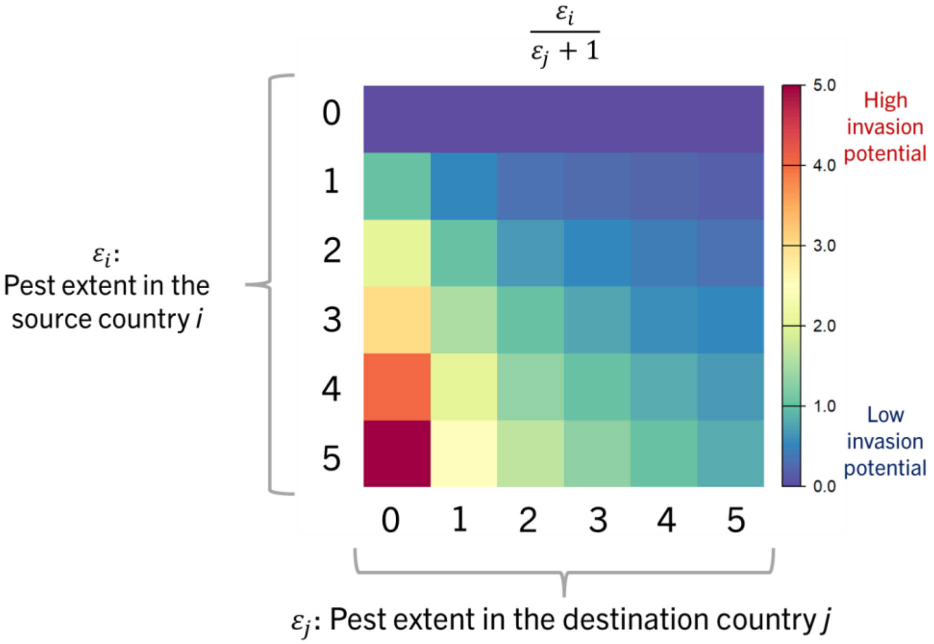


In general, this country-country formulation accounting only for pest extents allows us to consider the potential importance of introduction and re-introduction of a pest in a country $i$ relative to the pest extent in the country $j$.

- *Scenario 1.* One can expect that the larger the pest extent in the exporting country $j$, the greater the risk of pest introduction in the importing country $i$ if the pest is not yet present in country $i$. This hypothetical scenario is illustrated below. In this scenario, one can expect higher chances of unintentionally picking up or sampling propagule from the exporting country if the pest is widespread in the source country rather than geographically localized. The idea of 'picking up or sampling' here is an analogy to the quantity of possibly infected or contaminated crop commodities in the source country that is traded internationally.

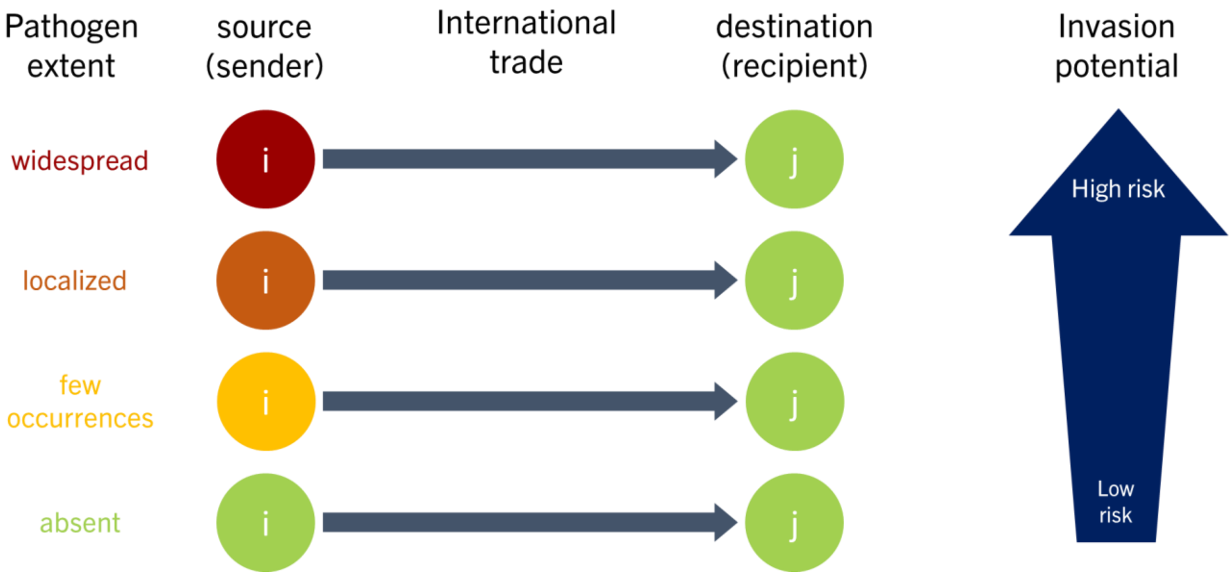


- *Scenario 2.* Likewise, the potential for pest re-introduction (or repeated introduction) in the importing country increases monotonically with larger pest extents in the exporting or source country. In this scenario, the pest is already present in both countries (scenario 2), and exporting crop commodities from a source country where the pest is widespread would allow a pest to be re-introduced more likely, more frequently or in greater quantities than from a source country where the pest is localized. Considering potential re-introductions is important because these events increase the likelihood that new genetic material of a pest species is potentially introduced and mixed with local populations in the destination country. For example, re-introduction is a particularly important concern for *Rastonia solanacearum* as this species comprises multiple sequevars, races, and biovars. Likewise, the strain of *Phthorimaea absoluta* introduced in Europe is likely clonal, but other strains from South America might pose a re-introduction risk worldwide.
- *Scenario 3.* This formulation ensures that there are always higher chances of moving pest propagule from source countries where the pest extent is greater (scenario 1 and 2), if pest movement is mediated by the international trade of crop commodities. If the pest is widespread in both countries (scenario 3), the potential movement of the pest from one country to another would be highly likely but is comparably not as important as if the pest was initially introduced in a pest-free country.

- *Scenario 4.* We finally assume that if the pest extent in the exporting country is zero (i.e., the pest is absent), the potential for pest introduction or re-introduction in the importing country is minimal but still potentially nonzero because of the possibility that some countries are re-exporting commodities from countries where the pest is present. We need to account for this idea of minimum baseline risk of pest introduction or re-introduction [43], even in the absence of the target pest in exporting countries.

To account for the above assumptions in our invasion potential analysis, we used a modified version of the above formulation in pest extents between countries as follows:

$$\tau_{i \to j} \propto \frac{\mathrm{A}^{\left(\frac{\varepsilon_i}{\varepsilon_j + 1}\right)}}{\mathrm{A}^5} \tag{1}$$

There are multiple beneficial features of this formulation of how invasion potential ($\tau_{i \to j}$) is proportional to a function of a positive scaling parameter (A) and pest extent in country $i$ ($\varepsilon_i$).

- The maximum invasion potential index is always 1 when the pest is widespread in the source country and is absent in the destination country (the worst-case scenario). The denominator in the function (i.e., $\mathrm{A}^5$) helps to keep invasion potential at a maximum value of 1.
- If there is international trade of commodities, the minimum invasion potential is not zero but a small value that depends on the base value of $\eta$. For example, the minimum invasion potential is 0.03125 if A = 2, $\varepsilon_i$ = 0, and $\varepsilon_j$= 0; and this small value indicates that pest introduction in the destination country is 32 times less likely than the maximum invasion potential. This formulation provides a risk-averse perspective, which is a key point considering the possibility of re-exported commodities transporting pests inadvertently.
- More importantly, this formulation allows us to conceptualize invasion potential as a non-linear function of pest extent, accounting for the typical exponential growth of a pest population in a host landscape. In this formulation, the base $n$ determines the steepness of the relationship between invasion potential and pest extent. This formulation of invasion potential based on pest extents is flexible in adjusting the base $n$ to any value, but we set it arbitrarily to 2 in our analysis. The plot below compares two values for the steepness of this formulation in pest extents ($\mathrm{A} = e$ or 2.71 in the left panel versus $\mathrm{A} = 2$ in the right panel). Note that different values in pest extents (x-axis) are needed to obtain the same levels of invasion potential (y-axis = 0.2).

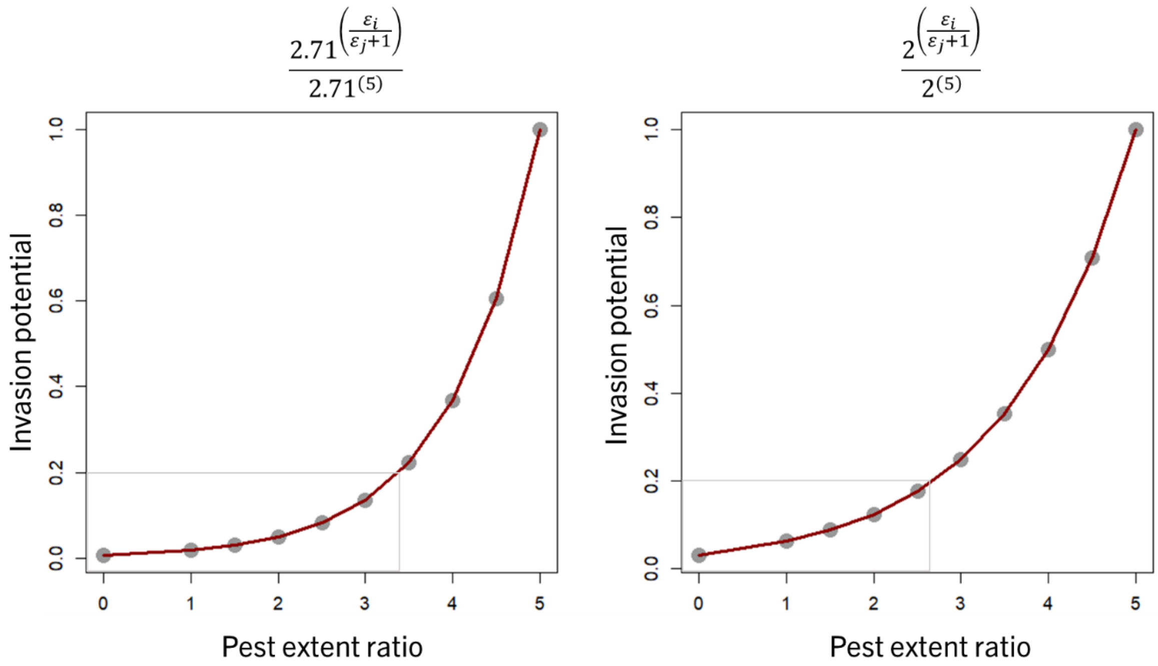


Notes on data sources: Currently available databases providing information on the countrywide extent of pests include the downloadable distribution lists in CABI Compendium (for example, https://doi.org/10.1079/cabicompendium.45009 for *Ralstonia solanacearum*), PlantwisePlus Knowledge Bank (https://plantwiseplusknowledgebank.org/; for example, https://doi.org/10.1079/pwkb.species.45009 for *Ralstonia solanacearum*), and EPPO Global Database (https://gd.eppo.int/; for example, https://gd.eppo.int/taxon/RALSSL/distribution for *Ralstonia solanacearum*). Many distribution lists require updates or to be created *de novo* based on published literature or unpublished expert knowledge. It is also important to note that non-reported pests do not always mean pest absence.

iii) Invasion potential as a function of international trade of commodities

The assumptions about invasion potential based on pest extent as suggested above are valid for comparison if trade between any pair of countries is constant. To incorporate the potential role of international trade on invasion potential, we assumed that the potential for pest introduction or re-introduction in a country is nonlinearly proportional to the amount of trade from the country $i$ to the country $j$. We propose three mathematical formulas that satisfy this expected nonlinear proportionality between trade and invasion potential, where we explain the context for these formulas below.

$$\tau_{i\to j} \propto \left(\frac{t_{\mathrm{M},i\to j}}{\max(t_{\mathrm{M},i\to j})}\right)^{\frac{1}{m}} + \left(\frac{t_{\mu,i\to j}}{\max(t_{\mu,i\to j})}\right)^{\frac{1}{m}}, \quad (2)$$

$$\tau_{i\to j} \propto \tanh\left(m * \frac{t_{\mathrm{M},i\to j}}{mean(t_{\mathrm{M},i\to j})}\right) + \tanh\left(m * \frac{t_{\mu,i\to j}}{mean(t_{\mu,i\to j})}\right), \text{ or} \quad (3)$$

$$\tau_{i\to j} \propto \frac{t_{\mathrm{M},i\to j}}{t_{\mathrm{M},i\to j}+mean(t_{\mathrm{M},i\to j})} + \frac{t_{\mu,i\to j}}{t_{\mu,i\to j}+mean(t_{\mu,i\to j})} \quad (4)$$

where $t_{\mathrm{M},i\to j}$ is the international trade volume of 'high-risk' commodities (that is, commodities playing a major role in the dispersal of a pest species), $t_{\mu,i\to j}$ is the international trade volume of 'low-risk' commodities (that is, commodities playing a minor role in the dispersal of a pest species), $m$ is a positive constant value, and max () (or $mean()$) are functions to calculate the

maximum (or mean) international trade volume across all international trade activities of a crop commodity.
In general, these three trade-invasion functions have the following features in common:

- They allow us to assume a high invasion potential with high volumes of international trade of a crop commodity. This is exemplified in a simple illustration below, where the pest extent in different exporting countries is the same, the pest is absent in the importing country, but there are different levels of export from exporting countries to the importing country.

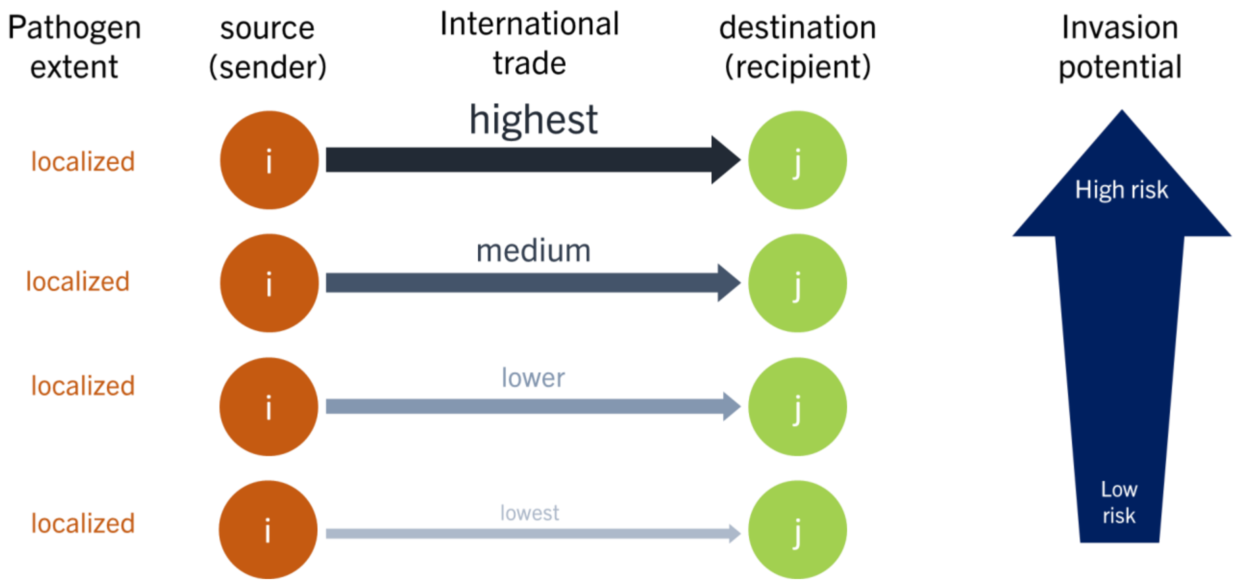


- These functions allow a non-linear association between trade volume and invasion potential (see figure below). This shape of invasion potential as a function of international trade has been suggested by some previous studies for pathogen and arthropod invasions [44-46]. This shape is expected because, historically, there has been a linear increase in the number of pathogen or insect invasions, while there has been an exponential growth in international trade [45,46]. We can think of trade volume as the effort size when sampling, where with larger sample sizes (or higher trade volumes) the chances for invasion accumulates until it reaches a saturation point [44]. In other words, we can think of trade volume as proportional to the number of repeated trials for the event of spread to occur.
- In other words, these nonlinear relationships between trade and potential pest movement capture canonical saturation expectations. The first function is a modified formulation of the log-log species-area model [44,47] and specifically captures a relationship based on power law models. The second function has been used to express saturation processes [48]. The third function is a modified formulation of the Michaelis-Menten model for the species-area model [44].

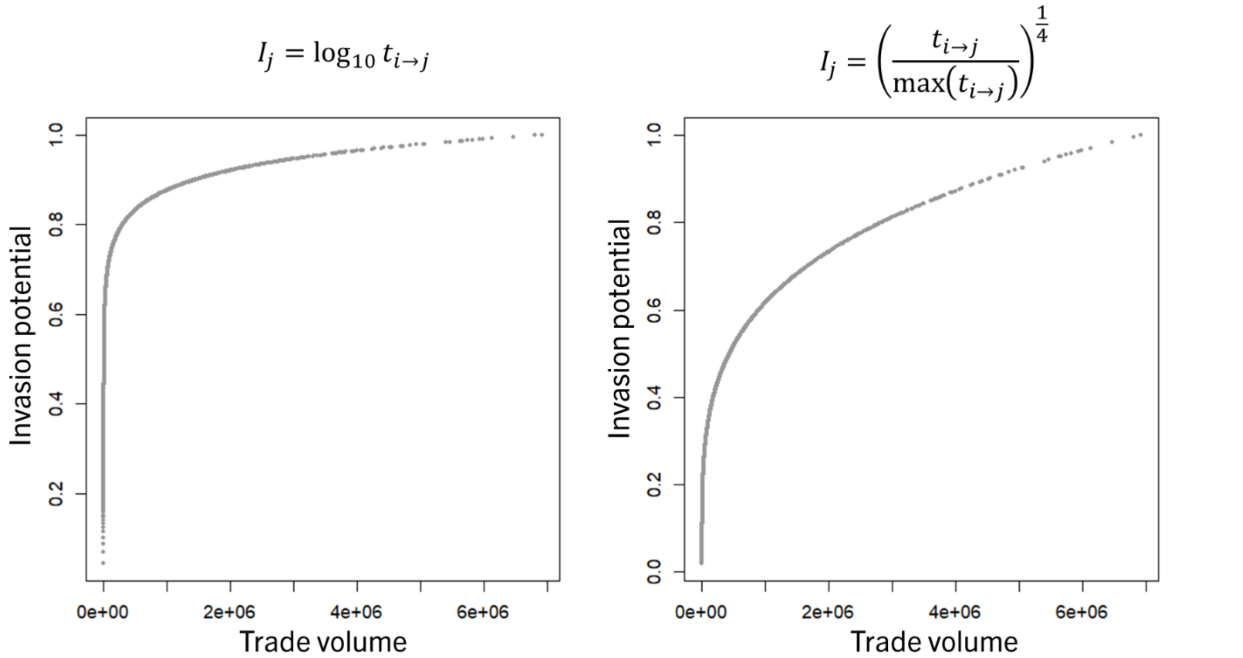


- They allow scaling invasion potential between 0 and 1. The shape of these functions can be adjusted by changing the values of the constant $m$. In our analysis, we used the first function with $m$ set at 4.

Trade data sources: (1) The FAOSTAT database provides a detailed trade matrix (https://www.fao.org/faostat/en/#data/TM) for the annual international trade of many commodity categories, reported by either the importing country (import quantity) or the exporting country (export quantity). We prefer reports on trade quantity (tons) rather than trade value (US$) to avoid price inflation issues. Since some countries lack trade statistics on import or export reports, we calculated the mean of international trade quantity across these two report types to include all possible trade activities between any pair of countries available in this database. Importantly, the designation of countries into 'reporting' and 'partner' categories in the FAO database should be switched in one report type to correctly keep the directionality of international trade when calculating the mean international trade. (2) The World Trade Organization (https://stats.wto.org/) provides a comparable database for annual bilateral imports value (US$) for many commodity categories. (3) The United Nations Comtrade Database (https://comtradeplus.un.org/) provides annual and monthly trade statistics, based on imports, exports, re-imports, or re-exports. Similarly as above, we prefer reports on trade quantity (kg) rather than trade value (US$) to avoid price inflation issues. An edgelist of international trade for specific commodities can be constructed with the csv files downloaded from any of above trade databases or a consensus across all databases. (4) Volza (https://www.volza.com/) provides historical trade statistics with more detailed commodity descriptions, including planting material and seed trade. However, this data source is publicly available for the most recent year's report only.

Notes on trade data caveats: (1) Not all databases provide specific information on the international trade of crop-specific planting material or seed, which is often the main pathway of spread for many plant pathogens and pests [46]. (2) Informal international trade of agricultural commodities (including planting materials) may not be reported in any of these databases, but these trade activities may pose a high risk for the spread of pathogens and pests. (3) The likelihood of commodity categories carrying or transporting a species is pathogen or pest-specific and depends on multiple factors. For example, frozen or heat-treated products may be less likely to transport pathogen inoculum or pest propagule than fresh products. Fenn-Moltu, et al. [49] provided an extensive dataset based on border interceptions to identify commodities associated with insect movement through international trade. There is not a comparable dataset and systematic analysis for pathogen-commodity associations published in a scientific journal. EUROPHYT (https://food.ec.europa.eu/plants/plant-health-and-biosecurity/europhyt/interceptions_en; https://food.ec.europa.eu/plants/plant-health-and-biosecurity/europhyt_en) provides monthly reports on interceptions of harmful (including pathogens) organisms in imported plants and other commodities. The EPPO Global Database also provides non-compliance reports for European countries (e.g., https://gd.eppo.int/reporting/article-7329), which include pest name (including pathogens), name of plant species consigned, type of commodity, country of origin, and country of destination based on interceptions. Expert knowledge can also be a source of unpublished or undocumented information regarding the associations between trade commodities and pest species. (4) Some commodity categories differ in resolution of classification or availability among trade databases. (5) Seasonality in the international trade of commodities is also important to understand the potential temporal dynamics in pest spread through the movement of commodities (e.g., roses are important on February 14) [50]. The UN Comtrade is the only database with monthly reports on commodity trade and Volza provides specific dates of trade activities but this information is not publicly available. (6) Including trade reports covering multiple years in the analysis is important due to time lags in pest establishment, spread and discovery [46,51]. A pest introduced through international commodity trade today may be discovered some years in the future. However, determining an effective time frame or the number of years to include in a trade analysis is challenging for most invasive species. (7) While total volume of international commodity trade provides a general picture of the possible introduction of a pest species (a component of propagule pressure), the frequency of trade may also indicate potential multiple (re)introductions (frequency of propagule pressure). Considering the number of trade activities is an alternative or complementary approach to trade volume. In the future, we should build databases and modelling approaches that help us understand quantitatively the probability, magnitude and duration of pest movement through international trade of commodities.

iv) Invasion potential as a function of host availability

A third aspect, the host population present in each pair of trading countries, is key for understanding pest invasion success for two reasons.

- The idea behind inclusion of this risk factor is that the size of the host population in the exporting country $i$ is a strong determinant of the invasion potential, which is needed to sustain a pest population. Thus, exporting countries having the same level of pest extent and the same level of export but different levels of host population ($\omega_i$) would increase the chances of 'sampling and exporting' a pest in countries with larger host populations than in smaller ones. Another motivation is that when we refer to a pest as widespread, it is likely that larger host populations can sustain larger pest populations compared to smaller host populations.

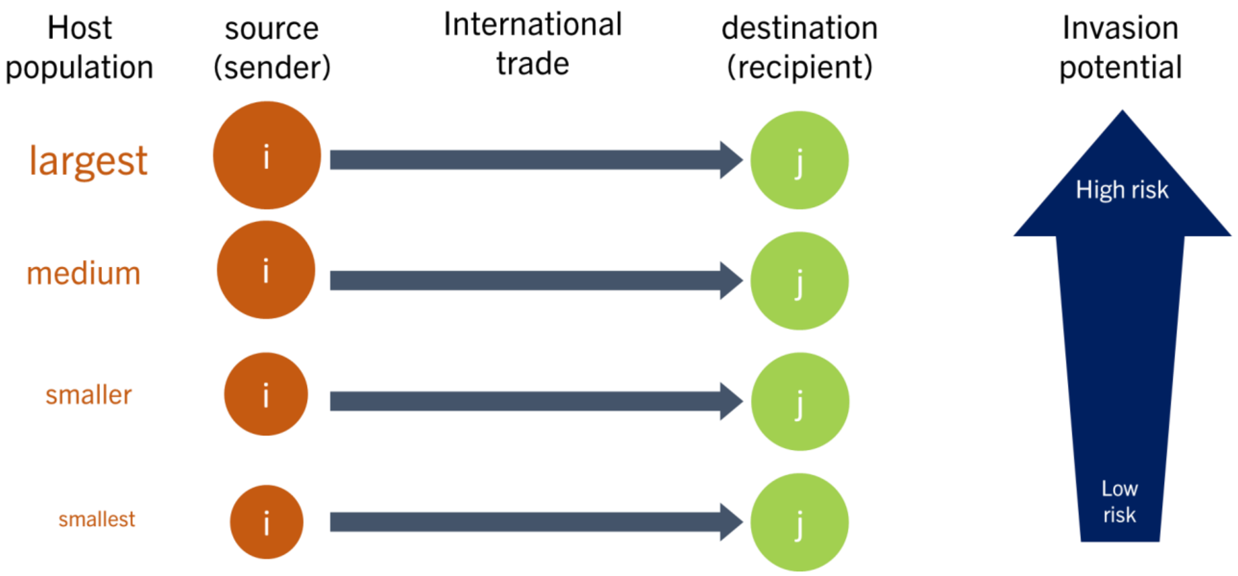


- Likewise, we hypothesize that the introduction or re-introduction of a pest in an importing country increases with larger host populations in the importing country ($h_j$).

Since previous studies indicate that the relationship between invasion potential and host population is logarithmic at small spatial scales [52,53], we incorporated a logarithmic function of host for quantifying invasion risk at large scales.

$$\tau_{i\to j} \propto \left(\frac{log(\Omega_i)}{\max(log(\Omega_i))} + \lambda_i\right) * \left(\frac{log(\Omega_j)}{\max(log(\Omega_j))} + \lambda_j\right), \text{ or} \quad (5)$$

$$\tau_{i\to j} \propto [1\left(\frac{\Omega_i}{\max(\Omega_i)}\right)^k + \lambda_i] * [1\left(\frac{\Omega_j}{\max(\Omega_j)}\right)^k + \lambda_j] \quad (6)$$

We add a parameter $\lambda$ (where our analyses use $\lambda_i = \lambda_j = \frac{1}{10}$) in this formulation to make all values of invasion potential greater than zero, and max() is the maximum host population available in any country in the world. One reason why invasion potential based on the host population might not be zero is the re-export of commodities. During an exploratory analysis of international trade, we noticed that some countries re-export crop commodities even if these countries do not produce these commodities (that is, these non-producing countries may not have available hosts within their territory). This re-exporting issue motivated us to include a minimum invasion potential in countries where hosts are unavailable ($\lambda$), because of the possibility that re-exporting countries are indirectly relocating crop commodities from pest sources. Again, we scaled invasion potential, so that the maximum value is one. We used equation (5) in our analysis.

Equation (6) is a central transmission-density function proposed by Hopkins, et al. [54]. Equation (6) is a modified formulation of $g = c(N/A)^k$, where $\frac{N}{A}$ is the host density, $c = 1$, and $k \in (0,1)$. In the theory of island biogeography, a similar formulation is proposed for the species-area curve $S = CA^z$, where S is the number of species of a given taxon found on a region and A is the area of the region, $k$ ranges between 0.12 and 0.17.

- Notes on host data sources: FAOSTAT (https://www.fao.org/faostat/en/#data/QCL) provides information about the harvested area of crop species at the national level. We used the harvested area of the crop species affected by the target pathogen as proxy for host availability within a country. Other data sources such as CROPGRIDs [55], MAPSPAM (https://mapspam.info/), and EARTHSTAT [56] are available for the geographic distribution of crop species and the Global Biodiversity Information Facility (GBIF; https://www.gbif.org/) for wild plant species. Information about plant species' distribution from these data sources needs to be aggregated at the country level for use in the pest introduction analysis, except for FAOSTAT.

- Notes on host data caveats. For pests with multiple plant host species, the role of host species during epidemics or pest invasion may be different, requiring consideration of the availability of primary ($H$) and secondary ($h$) hosts in the country. We can incorporate this potential role differences by assuming $\omega_i \propto \ln(AH_i + ah_i)$, where $H_i$ is the total abundance of all primary, major, or main hosts species of the targeted pest in country $i$ and $h_i$ is the total abundance of all secondary, minor, or alternative hosts species of the targeted pest in country $i$. The weights in our analyses allowed us to emphasize the importance of major hosts, where pests are more likely to survive and be transported in commodities. We arbitrarily set $A = 2$ and $a = 1$ in our analyses. Another important consideration for invasion potential is the seasonality in susceptible host availability. Pest invasion potential also depends on the geographic distribution of tolerant and resistant hosts at the intraspecies level. These two later factors are not considered in our analysis.

v) Invasion potential as a function of available biosecurity measures

Finally, we included the idea that the reported biosecurity level of the exporting country ($\beta_i$) and the biosecurity level of importing country ($\beta_j$) are inversely related to invasion potential in our analysis.

$$\tau_{i \to j} \propto \frac{1}{\beta_i * \beta_j} \quad (7)$$

We assumed that greater, more diverse, and more intense biosecurity efforts reduce invasion potential through international trade of agricultural commodities. For example, in 2023, the United States through a federal order imposed import requirements for tomato leafminer (*Phthorimaea absoluta*) hosts from infested countries. In this case, we classify countries as whether they are required to meet import requirements or not when exporting commodities (such as tomato fruits) to the United States. For the introduction potential analysis in the main paper, we set $\beta_i = 10$ if there was any type of pest-specific biosecurity measure imposed on traded commodities, and $\beta_i = 1$ otherwise. Risk analysts can modify these biosecurity parameter values ($\beta_i$ or $\beta_j$) as needed.

More generally, our analysis incorporates whether countries implement biosecurity measures when trading commodities internationally, which preferentially would be specific to the target pest species or specific to an agricultural commodity. This type of biosecurity information can be the implementation or adoption of International Standards for Phytosanitary Measures (ISPMs; https://www.ippc.int/en/core-activities/standards-setting/ispms/), which is reported by the International Plant Protection Convention (IPPC). For example, ISPM15 is crucial to prevent the movement of pests associated with wood packing material in international trade, and IPPC provides a list of countries implementing ISPM15 (https://www.ippc.int/en/countries/all/ispm15/). In the absence of a biosecurity measure specific to a pest species or agricultural commodity, an alternative approach is to consider general measures of countries' efforts in managing invasive species [e.g., Montgomery, et al. [40]]. A candidate measure for this alternative approach is the proactive and active response capacities of countries provided by Early, et al. [57], which account for multiple aspects of national responses to invasive species.

vi) Proportional relationships of invasion potential

Chapman, et al. [39] provided a multi-species pest invasion analysis for international trade of four broad categories of agricultural commodities (live plants, forest products, fruit and vegetables, and seeds) in Europe and the Mediterranean region. Gottwald, et al. [4] provided an introduction risk analysis for Asiatic citrus canker, citrus black spot, and citrus Huanglongbing (HLB) in the United States, where possible pest introduction is mediated by the international movement of people. Montgomery, et al. [40] also provided an example for the potential spread of the planthopper spotted lanternfly through the global trade network of stone commodities. To our knowledge, a detailed approach to estimating potential pest introduction through international trade networks of crop-specific commodities globally is lacking and needed. Available approaches though have not been applied to understand the possible introduction of plant pathogens.

Here our objective was to provide and apply such a detailed network-based modelling approach. In this modeling approach, we explicitly considered five major ecological, epidemiological, or biological factors for the potential introduction of pest species in a country (equation 8): crop-specific international trade ($t$), pest extent ($\varepsilon$), pest discovery duration ($\delta$), host availability ($h$), and biosecurity regulations ($\beta$). In a global trade network, the relative likelihood of accidentally introducing a pest species through commodity trade in an importing country ($\tau_j$) is proportional to the invasion potential posed by importing from a certain number ($z$) of exporting countries.

$$\tau_{i\to j} \propto \frac{A^{\left(\frac{\varepsilon_i}{\varepsilon_j+1}\right)}}{A^{(5)}} * \frac{1}{\beta_i * \beta_j} * \left(\frac{t_{i\to j}}{\max(t_{i\to j})}\right)^{\frac{1}{m}} * \left(\lambda_i + \frac{log(\Omega_i)}{\max(log(\Omega_i))}\right) * \left(\lambda_j + \frac{log(\Omega_j)}{\max(log(\Omega_j))}\right) \quad (8)$$

We assume that $P(\tau_{i\to j})$ represents the relative likelihood of pest movement from country $i$ to $j$. If a destination or importing country is trading only with one source or exporting country, $1 - P(\tau_{i\to j_1})$ is the relative likelihood that trading agricultural commodities from country $i$ to $j$ does not transport or carry pest propagule. If the importing country is trading with two exporting countries, $(1 - P(\tau_{i\to j_1}))(1 - P(\tau_{i\to j_2}))$ is the joint relative likelihood that neither of two exporting countries $i$ introduce the target pest species into country $j$ by the trade of agricultural commodities. We assume that the events of introduction from these two countries are independent of each other. More broadly, under the same assumption of independence, the joint relative likelihood that none exporting countries introduce the pest species into a target importing country is $\prod_{k=1}^{z}(1 - P(\tau_{i\to j_k}))$, where $z$ is the number of countries exporting agricultural commodities to country $j$. Finally, the joint relative likelihood that the pest is introduced into a country from any exporting country is $I_j$, where

$$I_j \propto 1 - \prod_{i=1}^{z}(1 - P(\tau_{i\to j_k})) \quad (9)$$

Note that our formulation of $\tau_{i\to j}$ explicitly incorporates the three major components of species dispersal as envisioned in movement ecology or dispersal biology: source processes (departure or emigration), relocation processes (transience or transfer), and destination processes (settlement or immigration)[58-61]. Among the risk factors included in equation (8), international trade of commodities is most likely associated with relocation processes, while the other factors are more likely to influence source or destination processes in a global landscape.

vii) Combining ports, cities and trade connectivity

Because international trade is only available at country resolution, we allocated each grid in the global accessibility to cities and ports as belonging to its corresponding country. For each country, we used $I_j$. We then disaggregated $I_j$ using accessibility to ports ($I_p$) and cities ($I_c$) to approximate initial entry points or final destinations into countries. Our final index of pest introduction potential due to trade (or $\Delta_t$) is proportional to port accessibility within a country ($I_p$) and the country-level introduction risk when importing commodities ($I_j$):

$$\Delta_t \propto I_p \times I_j \tag{10}$$

The assumptions of combining port or city accessibility with $I_j$ are provided in the main text of this paper.

viii) Future considerations on the introduction potential based on international trade

Our analytical framework for assessing pest introduction potential, as detailed above, originated from a growing interest in applying network analysis to enhance the understanding of biological invasions [62]. Below, we briefly mention potential future avenues to improve this analytical framework if the required geographic data becomes available.

- In equation (8), each geographic risk factor is equally weighted and is equally likely to influence invasion potential. In some cases, geographic risk factors might exhibit different levels of importance in invasion potential. Thus, a generalized formulation of equation (8) is equation (11):

$$\tau_{i\to j} \propto x_1 \ln\left[\frac{A^{\left(\frac{\varepsilon_i}{\varepsilon_j+1}\right)}}{A^{(5)}}\right] + x_2 \ln\left[\left(\frac{t_{i\to j}}{\max(t_{i\to j})}\right)^{\frac{1}{m}}\right] + x_3 \ln\left[\left(\frac{log(h_i)}{\max(log(h_i))} + \lambda_i\right) * \left(\frac{log(h_j)}{\max(log(h_j))} + \lambda_j\right)\right] + x_4 \ln\left[\frac{1}{\beta_i * \beta_j}\right] \tag{11}$$

  Where $x_1$,…, $x_4$ are specific weights assigned to each geographic risk factor. However, determining the specific importance of geographic risk factors for the introduction potential of a pest species is challenging.

- In their risk-based introduction analysis, Gottwald, et al. [4] considered infection duration as an important factor for estimating the relative plant pathogen strength in a source country. We expand this assumption about pest duration in a source country ($\delta_i$), where earlier pest detections may indicate that a pest could have had greater opportunities for spread within a country. We defined pest duration ($\delta_i$) as the difference between the current year and the year of earliest detection of the pest species in the source country. Including this assumption about pest duration is particularly important for pest species with a restricted geographic distribution but that continues expanding over other host areas.
- Our index of pest introduction potential (equation 10) does not incorporate the role of environmental factors [39], which are usually pest-specific conditions; stochasticity [40]; and time lags in species discovery [46].

- Our pest introduction potential model ignores the principle of competitive exclusion between pest species, other interspecific interactions such as the populations of natural enemies of pest [63], and the Allee effect that some pests may experience [64].

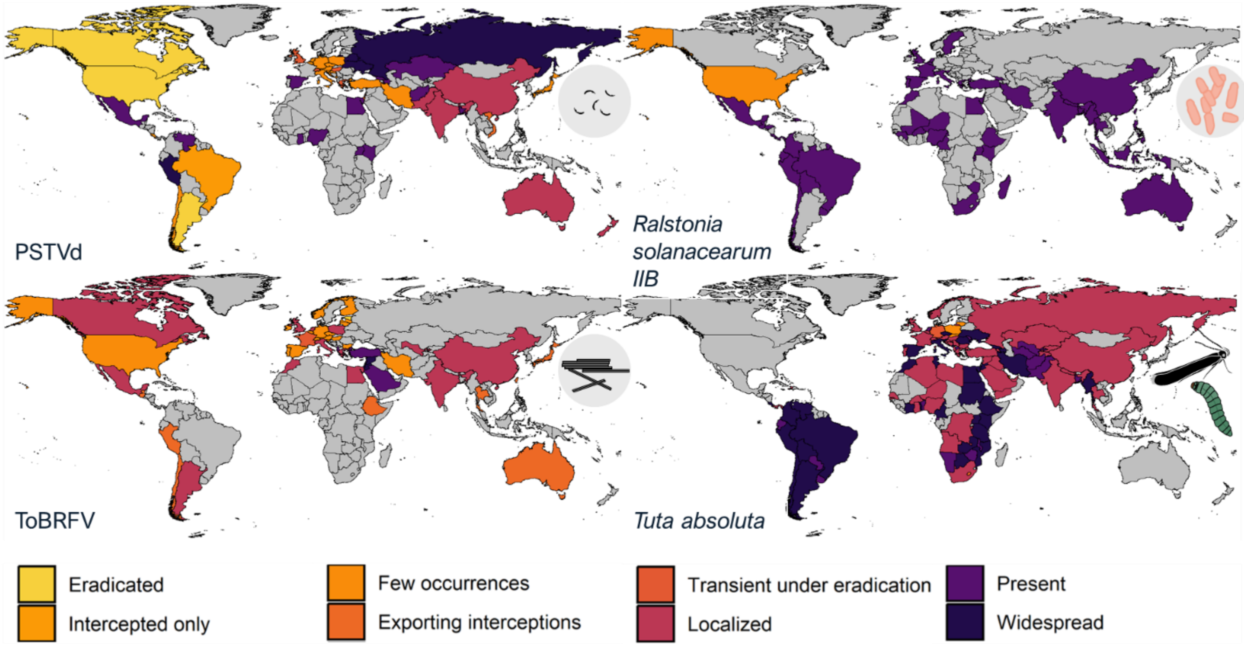


**Fig. S1.**
Potential pest sources for sentinel surveillance based on reported within-country distribution.

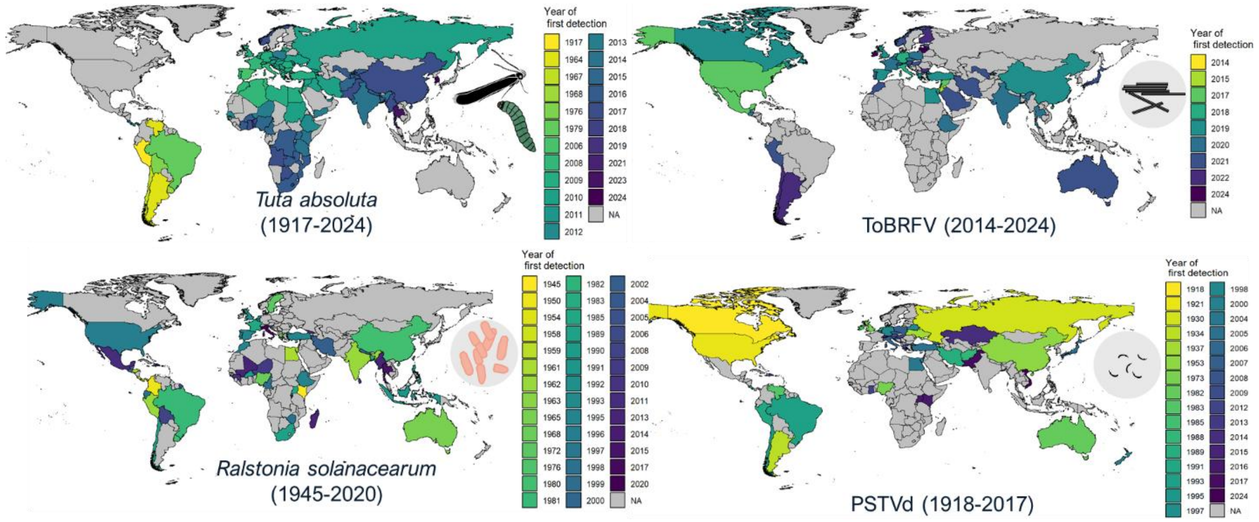


**Fig. S2.**

Geographic pest discovery reconstructed from earliest national observations.

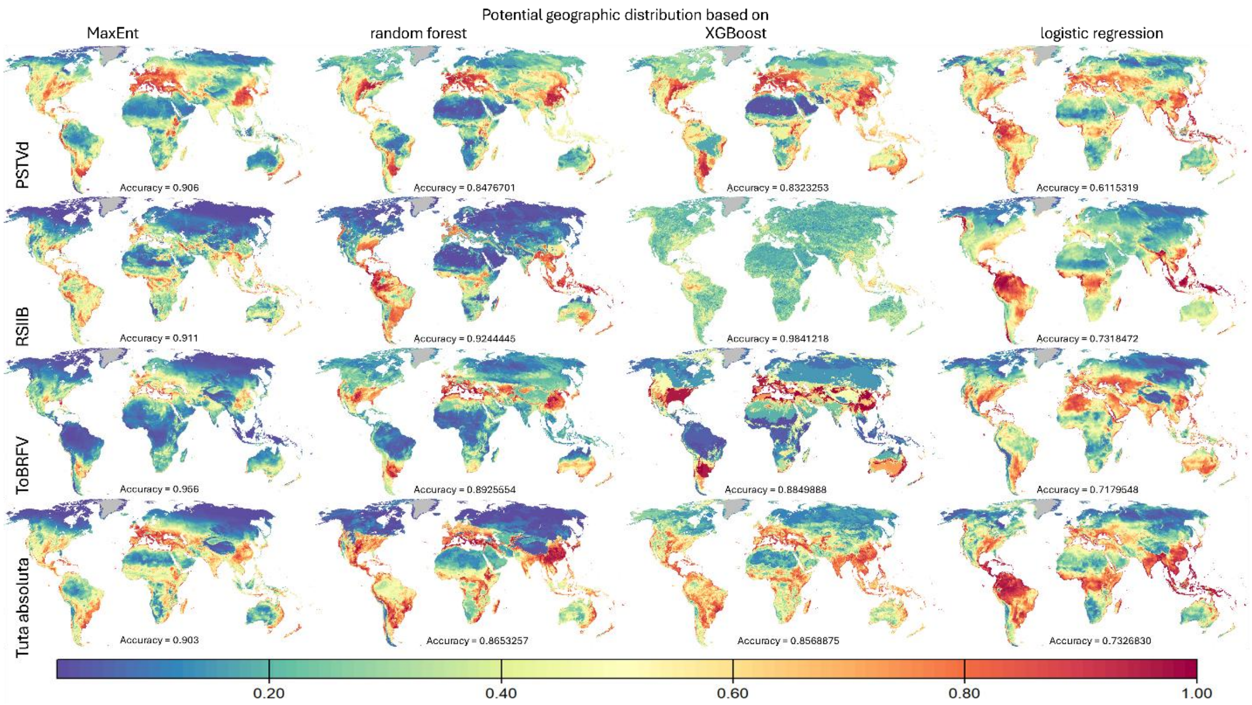


**Fig. S3.**

Predictions for pest presence based on four machine-learning algorithms.

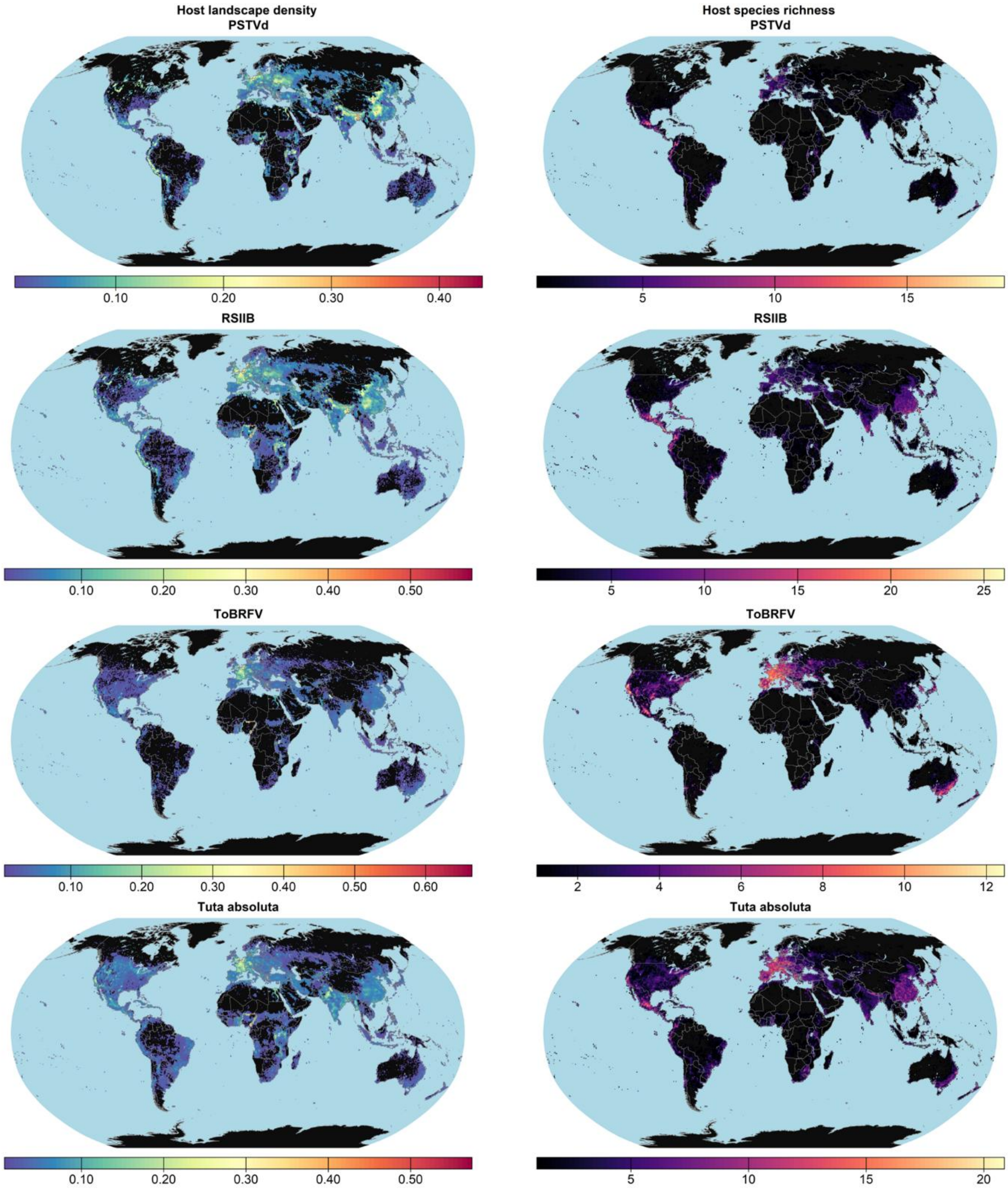


**Fig. S4.**

Geographic patterns in cumulative host density and host species richness for each target pest.

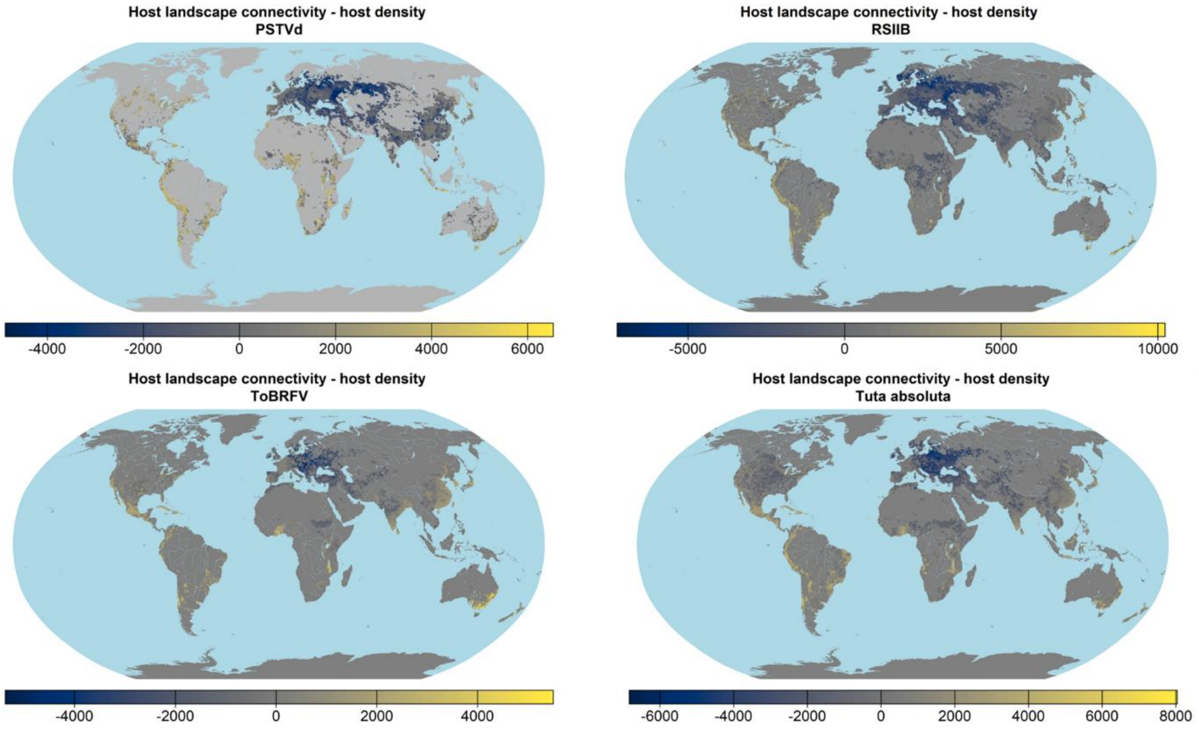


**Fig. S5.**
Maps of difference in ranks between mean host landscape connectivity and total host density.

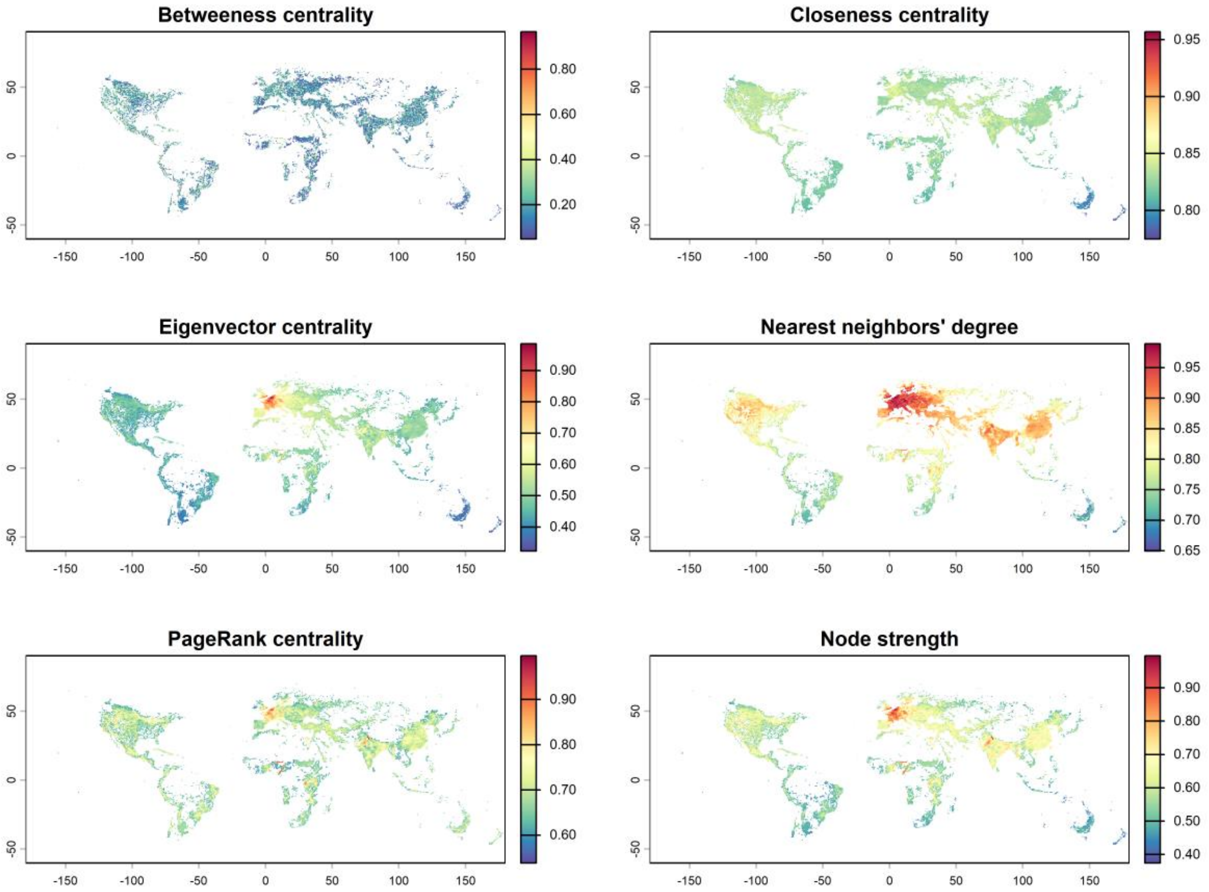


**Fig. S6.**
Maps of host landscape connectivity for *Phthorimaea absoluta* based on six network metrics.

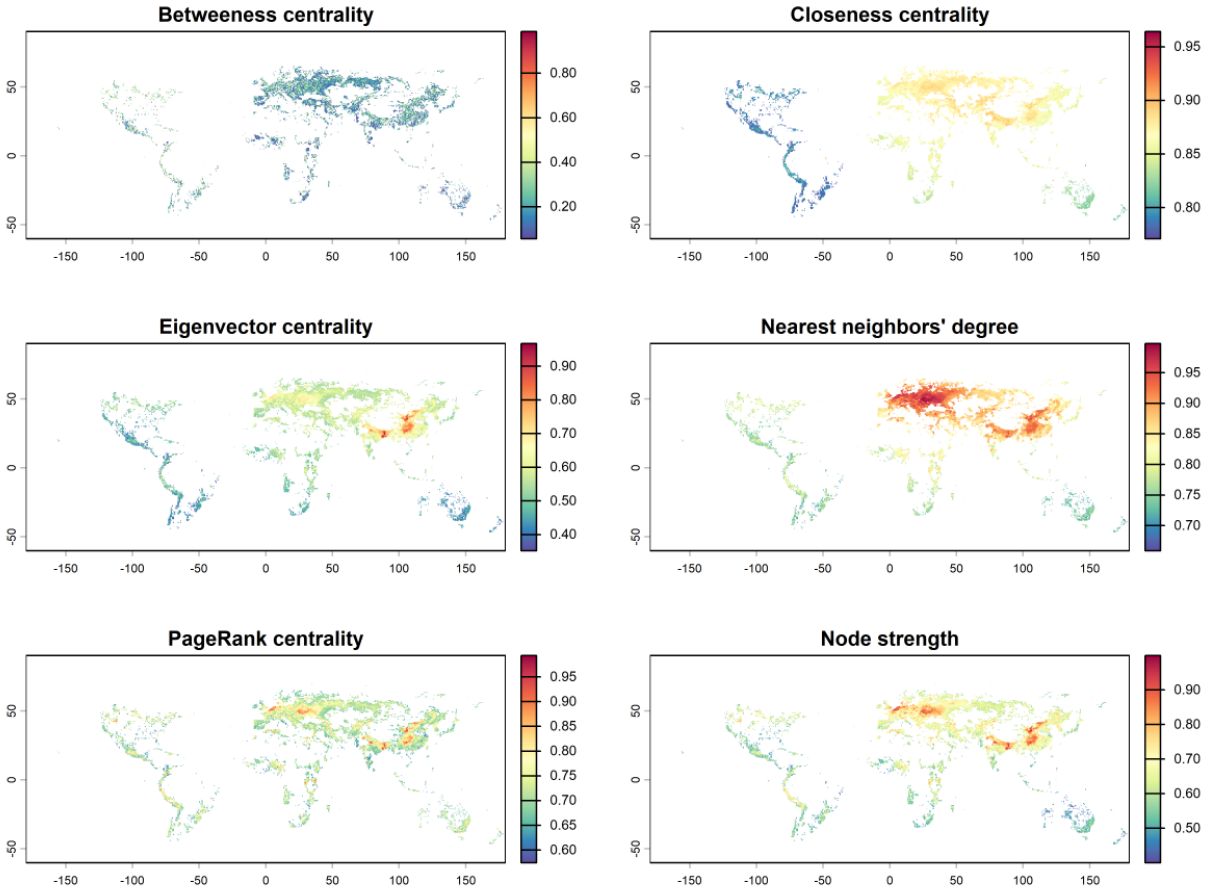


**Fig. S7.**
Maps of host landscape connectivity for potato spindle tuber viroid based on six network metrics.

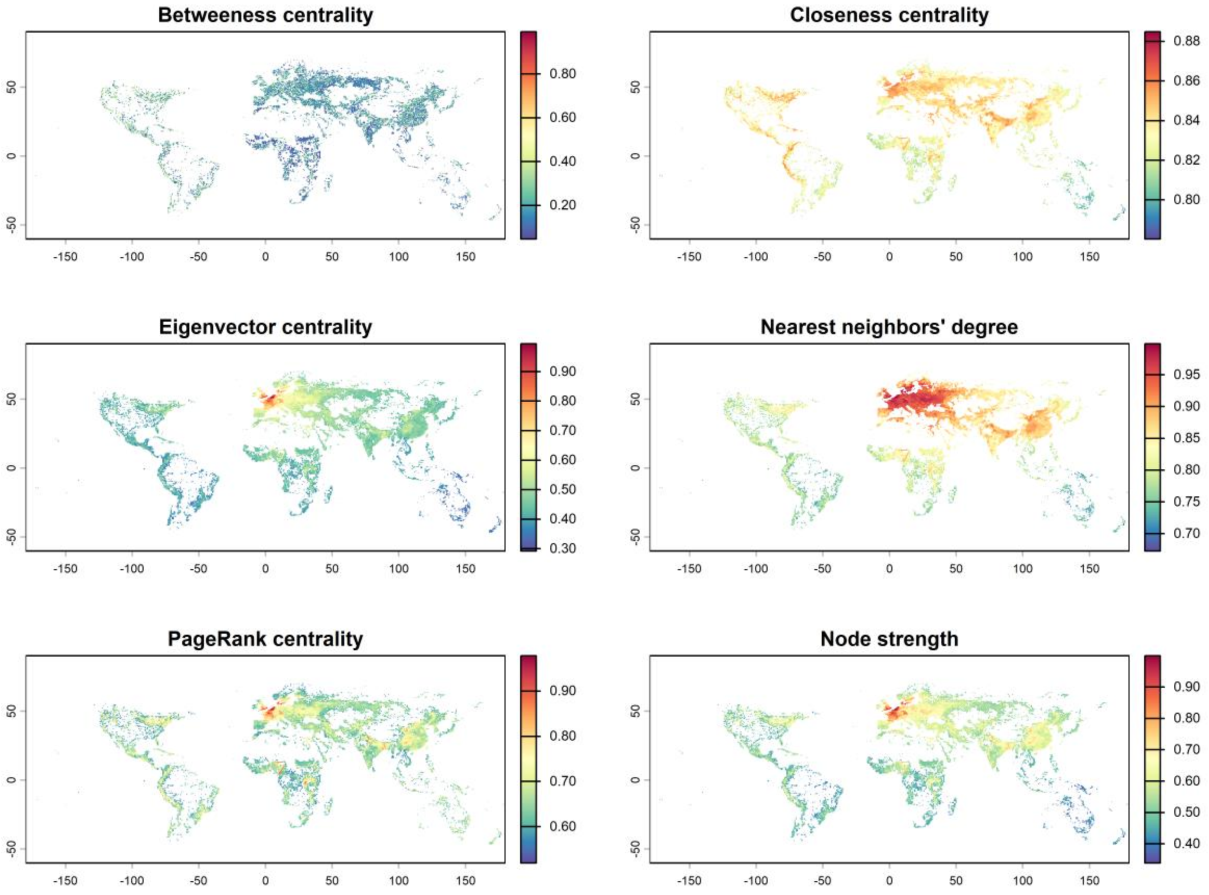


**Fig. S8.**
Maps of host landscape connectivity for *Ralstonia solanacearum* phylotype II-Bbased on six network metrics.

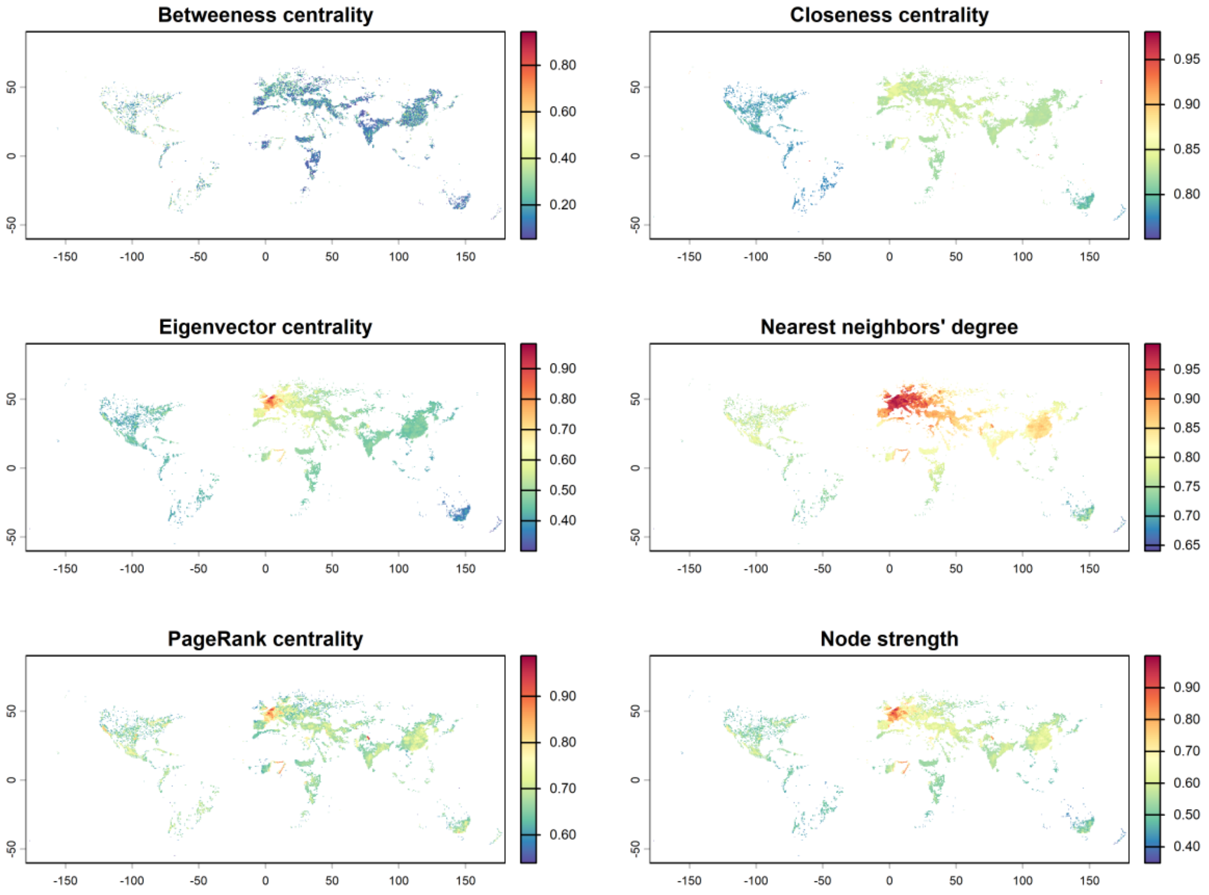


**Fig. S9.**
Maps of host landscape connectivity for tomato brown rugose fruit virus based on six network metrics.

**Table S2.**

Categories and weights assigned to global maps of accessibility to ports.

| Port ID | Port size | Number of ports | Model weight | Likelihood rank |
|---|---|---|---|---|
| **A** | Large | 160 | 0.35 | Highly likely |
| **B** | Medium | 361 | 0.3 | Mild likely |
| **C** | Small | 990 | 0.2 | Low likely |
| **D** | Very small | 2153 | 0.15 | Very low likely |
| **Total map** | All sizes | 3664 | 1 | Any likely |
| | Average accessibility = $\frac{0.35A+0.3B+0.2C+0.15D}{4}$ | | | Any likely |

**Table S3.**

Categories and weights assigned to global maps of accessibility to cities.

| Settlement class | Minimum population threshold | Maximum population threshold | Number of settlements | Population in settlements | Weight |
|---|---|---|---|---|---|
| **1** | 5,000,000 | 50,000,000 | 79 | 941,207,809 | ½ |
| **2** | 1,000,000 | 5,000,000 | 421 | 851,153,118 | 1/3 |
| **3** | 500,000 | 1,000,000 | 581 | 400,180,511 | ¼ |
| **4** | 200,000 | 500,000 | 2,096 | 630,823,940 | 1/5 |
| **5** | 100,000 | 200,000 | 3,694 | 515,557,120 | 1/6 |
| **6** | 50,000 | 100,000 | 6,973 | 484,166,417 | 1/7 |
| **7** | 20,000 | 50,000 | 20,457 | 628,095,955 | 1/8 |
| **8** | 10,000 | 20,000 | 29,286 | 410,631,333 | 1/9 |
| **9** | 5,000 | 10,000 | 45,795 | 322,797,326 | 1/10 |

**Table S4.**

Major host species of the target pest used in the multi-host connectivity analysis. Note that minor host species used in the analysis are not mentioned here.

| Target pest name | Abbreviation | Main host species |
|---|---|---|
| *Phthorimaea absoluta* | | Tomato (*Solanum lycopersicum*) |
| *Tomato brown rugose fruit virus* | ToBRFV | Tomato (*Solanum lycopersicum*)<br>Peppers (*Capsicum annuum*) |
| *Ralstonia solancearum* phylotype IIB1 | RSIIB1 | Bittersweet (*Solanum dulcamara*)<br>Potato (*Solanum tuberosum*)<br>Tomato (*Solanum lycopersicum*) |
| *Potato spindle tuber viroid* | PSTVd | Potato (*Solanum tuberosum*) |

**Table S5.**
List of agricultural commodities associated with each target pest and used as a dispersal pathway in the trade-mediated (re)introduction risk analysis.

| Target pest name | Commodity | Importance | Units | Data source |
|---|---|---|---|---|
| *Phthorimaea absoluta* | HS 0702.00 Tomatoes, fresh or chilled | Major (100%) | Tons | World Trade Organization (WTO) |
| *Tomato brown rugose fruit virus* | Tomato seeds | Major (100%) | Standard Qty | Volza |
| | HS 0702.00 Tomatoes, fresh or chilled | Minor (50%) | Tons | World Trade Organization (WTO) |
| *Ralstonia solancearum* phylotype IIB1 | HS0701.10 Potatoes, seed | Major (100%) | Tons | World Trade Organization (WTO) |
| | Pelargonium, geranium plants, geranium seeds, or geranium for sowing | Major (100%) | Standard Qty | Volza |
| *Potato spindle tuber viroid* | HS0701.10 Potatoes, seed | Major (100%) | Tons | World Trade Organization (WTO) |
| | Petunia flower, plants, seeds, and for sowing | Major (100%) | Standard Qty | Volza |
| | Capsicum seeds, chilli seeds, sweet, hot, green pepper seeds, red pepper seeds, charliston pepper seeds, sweet or bell pepper seeds, capsicum plants | Major (100%) | Standard Qty | Volza |

**Table S6.**
Model performance metrics for the multi-host connectivity analysis.

| **Pest name** | **Precision** | **A** | **B** | **Test statistic $D^+$** | **p** |
|---|---|---|---|---|---|
| Potato spindle tuber viroid | 0.827 | 0.201 | 0.173 | 0.26621 | 0.0001586 |
| *Ralstonia solanacearum* phylotype IIB | 0.867 | 0.192 | 0.138 | 0.32822 | 8.241e-13 |
| Tomato brown rugose fruit virus | 0.677 | 0.225 | 0.126 | 0.45953 | 1.412e-12 |
| *Phthorimaea absoluta* | 0.810 | 0.155 | 0.131 | 0.25709 | < 2.2e-16 |
| Average | 0.792 | 0.192 | 0.141 | | |

Precision: The ratio between the number of grid cells where the target pest has been reported present and the multi-host connectivity is nonzero (true positives) and the number of grid cells where the target pest has been reported present (true positives + false negatives). Note that our validation is based on presence-only data, so false positives and true negatives are not calculated.

A: The mean value in multi-host connectivity of grid cells where there are georeferenced observations of the reported presence of a pest species.

B: The mean value in multi-host connectivity across all grid cells, which were relevant for the multi-host connectivity analysis of each pest species.

**Table S7.**
Model performance metrics for the (re)introduction vulnerability analysis.

| Pest name | Precision | A | B | Test statistic $D^+$ | p |
|---|---|---|---|---|---|
| Potato spindle tuber viroid | 1 | 0.171 | 0.107 | 0.24615 | 0.2746 |
| *Ralstonia solanacearum* phylotype IIB | 1 | 0.078 | 0.100 | 0.08772 | 0.8037 |
| Tomato brown rugose fruit virus | 1 | 0.047 | 0.118 | 0.01449 | 0.9293 |
| *Phthorimaea absoluta* | 1 | 0.134 | 0.028 | 0.15607 | 0.5406 |
| Average | 1 | 0.094 | 0.087 | 0.06123 | 0.7296 |

Precision: Ratio between the number of countries where the pest has been intercepted on imported agricultural commodities and introduction risk was non-zero (true positives) and the number of countries where the pest has been intercepted on imported agricultural commodities (true positives + false negatives). Note that our validation is based on presence-only data, so false positives and true negatives are not calculated.

A: Mean value in (re)introduction risk in countries where the pest has been intercepted on imported agricultural commodities.

B: Mean value in (re)introduction risk in countries where the pest has not been intercepted on agricultural commodities.

**Data S1-12. (separate file)**

These datasets were used to construct models in GIRAF. See a separate file for Supplementary Data.
